# Spatial domains identification in spatial transcriptomics by domain knowledge-aware and subspace-enhanced graph contrastive learning

**DOI:** 10.1101/2024.05.09.593192

**Authors:** Yang Gui, Chao Li, Yan Xu

## Abstract

Spatial transcriptomics (ST) technologies have emerged as an effective tool to identify the spatial architecture of the tissue, facilitating a comprehensive understanding of organ function and tissue microenvironment. Spatial domain identification is the first and most critical step in ST data analysis, which requires thoughtful utilization of tissue microenvironment and morphological priors. To this end, we propose a graph contrastive learning framework, GRAS4T, which combines contrastive learning and subspace module to accurately distinguish different spatial domains by capturing tissue microenvironment through self-expressiveness of spots within the same domain. To uncover the pertinent features for spatial domain identification, GRAS4T employs a graph augmentation based on histological images prior, preserving information crucial for the clustering task. Experimental results on 8 ST datasets from 5 different platforms show that GRAS4T outperforms five state-of-the-art competing methods in spatial domain identification. Significantly, GRAS4T excels at separating distinct tissue structures and unveiling more detailed spatial domains. GRAS4T combines the advantages of subspace analysis and graph representation learning with extensibility, making it an ideal framework for ST domain identification.

## Introduction

The emergence of spatial genomics has revolutionized biological and medical research, facilitating an unprecedented understanding of functional arrangement. Knowledge of the relative locations of different cells in a tissue is critical for understanding tissue development because spatial information helps to link gene-coded tissue structural development to different functional domains. Deciphering spatial domains, i.e., identifying spatial spots with coherent gene expression and histology, is considered as a critical step in spatial transcriptomics analyses. It aims to assign spatial data into a series of meaningful clusters, where each cluster is considered as a spatial domain^1^. The successful identification of spatial domains usually relies on the effective utilization of gene expression data, spatial coordinates, and the corresponding tissue image^2, 3^, offering the potential to unveil inherent interactions and characterize the tissue microenvironment.

Several computational tools have been developed to decipher spatial domains. These methods can be classified as non-spatial- and spatial-based methods, depending on whether or not incorporate the spatial information. Among the non-spatial-based methods, classical clustering methods such as Louvain^4^, mclust^5^, and kmeans++^6^ were proposed. These methods neglect spatial information and histology, leading to inconsistent identification domains and reduce accuracy compared to spatial-based methods like BayesSpace^7^, Giotto^8^, and SC-MEB^9^, etc. In addition to the machine learning methods mentioned earlier, several deep learning-based methods^10–15^ have also been developed within the categories of spatial-based methods. Notably, there have been extensive research on methodologies based on Graph Neural Network (GNN). Given that spatial transcriptomics data include spatial coordinates naturally linking individual spots, the spatial domain identification challenge can be transformed into a node classification problem that the GNN has been proposed to solve. GNN-based methods convert the ST data into graph structures, enhancing data quality for downstream tasks by extracting deep graph structure information. Many GNN-based models employ autoencoder architecture or contrastive strategy to learn the representation of each spot. Particularly, the graph attention autoencoder framework STAGATE^14^ employs a cell type-aware attention mechanism to learn low-dimensional embeddings by integrating spatial information and gene expression profiles. DeepST^11^ identifies spatial domains based on denoising autoencoder and graph autoencoder. By integrating spatial local relations between gene expression, spatial location, and histological image, DeepST enhances gene expression and feeds it into a deep representation learning model to learn low-dimensional representation. The graph contrastive learning-based methods provide new perspectives into spatial transcriptomics data analysis. CCST, as proposed by Li et al.^12^, employs gene expression profiles and hybrid adjacency matrices based on cellular neighborhoods as inputs to a neural network that encodes cellular embeddings from spatial transcriptomics data. SpaceFlow^13^ makes the distance between learned potential embeddings mimic the distance between the gene expressions of a spot or cell, as well as the distance between spatial locations. This is achieved by adding a regularization term to maintain a level of consistency across these distances. conST^15^ uses three types of spatial transcriptomics data and takes into account the global, local, and context information of the graph. By leveraging this information, conST effectively learns the underlying graph structure of the data. While these methods take into account nearest neighbor relationships between spots, they fall short in adaptively distinguishing or identifying truly related spots. As a result, they struggle to precisely demarcate spatial domains. Moreover, most of these methods use domain-agnostic graph augmentations (DAGA) that do not consider spatial segmentation-related information, which leads to poor discriminative representation^16, 17^.

To address the challenges mentioned above, we propose GRAS4T, a domain knowledge-aware GRAph contraStive learning based on Subspace-enhanced for Spatial domain identification in Spatial Transcriptomic. GRAS4T initially incorporates a morphology prior-based graph augmentation module, which guides the connection and disconnection of edges within the spot-spot graph. The module constructs positive views that retain morphological information, enabling GNN to learn domain segmentation-related representation by way of contrast. For adaptive obtaining of real spot nearest neighbor information, GRAS4T incorporates the subspace module into the contrastive learning framework. Unlike the subspace model alone, GRAS4T uses latent feature space to obtain a self-expression matrix with better block diagonal properties^18^. With the self-expression matrix, the contrastive view contains realistic tissue microenvironment adaptively, thus obtaining more features relevant to the downstream clustering task. GRAS4T is applicable to various platforms (e.g., 10x Visium, Stereo-seq, MERFISH, etc.) and tissues (e.g., brain, breast, olfactory bulb, etc.), and is highly interpretable and extensible due to its combination of machine learning and deep learning.

## Materials and methods

### Data description

We collected eight sets, comprising 38 sections, of spatial transcriptomics datasets in this study. These datasets covered different platforms, tissues, sizes, and clusters (Supplementary Table S1). Four datasets were from the 10X Visium platform, encompassing various tissues and species: the human dorsolateral prefrontal cortex (DLPFC)^19^, human breast cancer, mouse brain anterior&posterior, and coronal mouse brain. The DLPFC dataset was obtained by spatially analyzing gene expression in two pairs of ‘spatial replicates’ sections from three independent adult donors. Each pair consisted of two consecutive tissue sections 10 μm apart, with the second pair located 300 μm behind the first pair. In total, there were 12 slices, with each slice containing from 3,460 to 4,789 spots and capturing 33,538 genes. Furthermore, the coronal section of the adult mouse brain only had one slice which contained 2,903 spots with 32,285 genes. The one slice of the human breast cancer dataset contained 3,798 spots with 36,601 genes. Lastly, the mouse brain anterior&posterior dataset^20^ contained 2,823 and 3,289 spots with 32,285 genes, respectively.

The remaining four datasets originated from various platforms and featured different spatial resolutions. Specifically, the human HER2-positive breast tumor (HER2+) dataset^21^ was generated using the spatial transcriptomics platforms^22^. The HER2+ dataset comprised tumor tissue sections from eight HER2-positive patients (A-H). Researchers collected a total of 36 samples, extracting either three or six sections from each tumor for analysis. In this dataset, each slice contained from 167 to 659 spots and captured 14,861 to 15,661 genes. The mouse visual cortex dataset was generated based on STARmap^23^, which contained 1,207 spots with 1,020 genes. The mouse olfactory bulb^24^ with a spatial resolution of 14 μm, was produced by Stereo-seq^25^, which contained 19,109 spots with 27,106 genes. Lastly, the mouse hypothalamus dataset was generated by MERFISH^26^, a high-throughput and highly multiplexed method based on *in situ* hybridization technology. We selected all slices from one mouse, with each slice containing from 4,787 to 6,154 spots and capturing 155 genes.

### Overall architecture of GRAS4T

GRAS4T uses the graph contrastive learning framework as its backbone, incorporating domain knowledge-aware graph augmentation and a subspace module to distinguish tissue boundaries. As illustrated in Fig. 1, the architecture of our GRAS4T contains four components, namely **data preprocessing, graph augmentation, domain feature extraction, and downstream task**. Given an input graph 𝒢 (defined in Supplementary section 2.1), it employs the well-designed graph augmentation techniques to obtain the positive pairs and uses the corruption function to obtain the negative pairs. These graphs serve as inputs to the GNN-based encoder to obtain low-dimensional representations. Next, the node representation obtained from the input graph 𝒢 is subsequently fed into the subspace module to get the self-expression matrix. Unlike the classic contrastive learning structure, GRAS4T adopts the self-expression matrix to obtain the reconstructed representation of positive views, which participates in the computation of the local-subspace contrastive loss. Furthermore, GRAS4T uses the decoder to recover the data matrix so that embedding retains sufficient gene expression-related information. The whole model is trained with local-global contrastive loss, local-subspace contrastive loss, and reconstruction loss. Finally, the node representation is applied for downstream tasks such as clustering. Detailed explanations of these modules will be elaborated in the subsequent sections.

**Figure 1.**
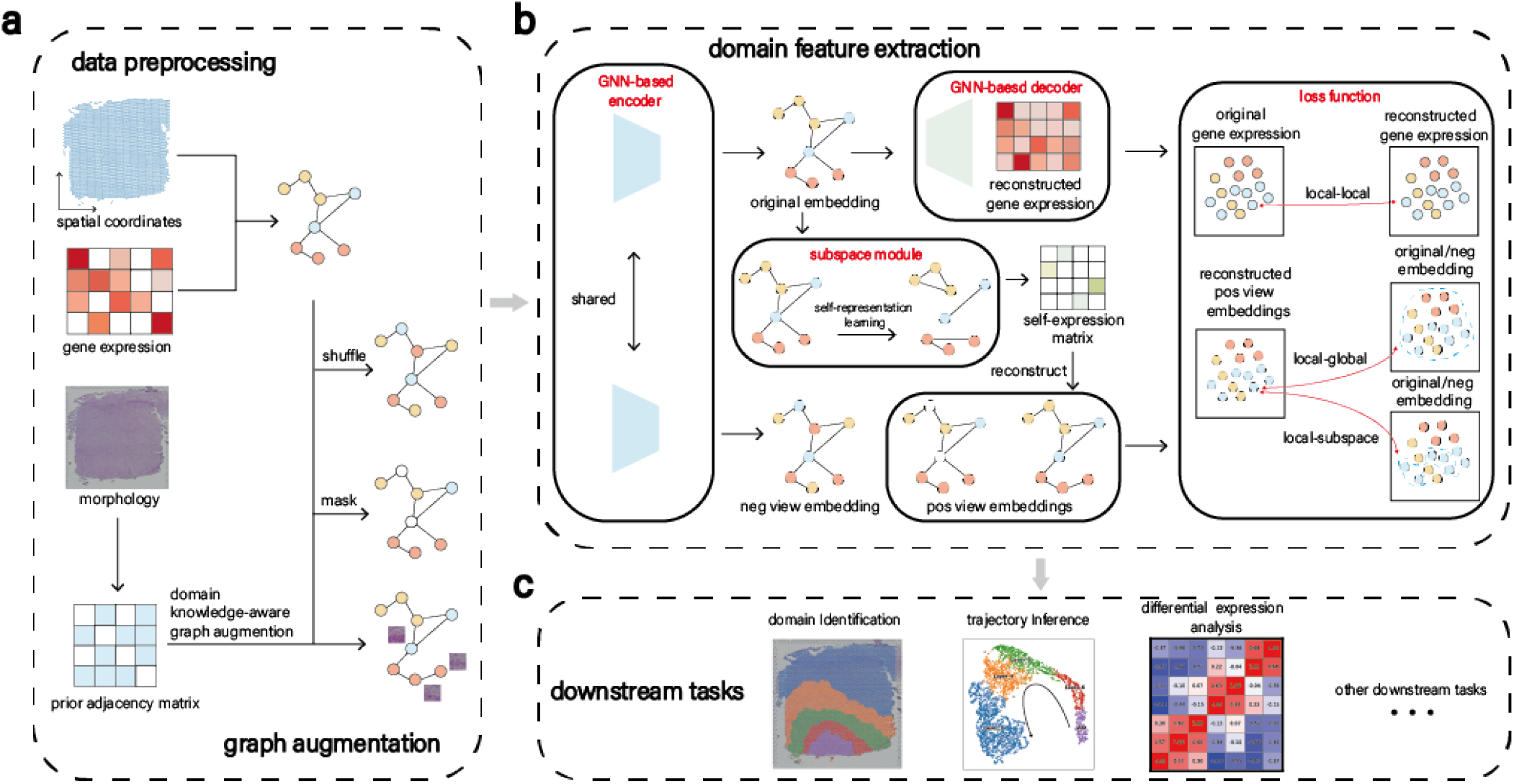
Overview of GRAS4T. (a) GRAS4T constructs graph 𝒢 from gene expression profiles and spatial locations, then uses mask and shuffle to augment it, creating positive and negative views. The prior adjacency matrix is derived from the hematoxylin and eosin (H&E) image, while the positive view is obtained by augmenting graph 𝒢 with the help of the prior adjacency matrix. Graph 𝒢 along with the three generated views will serve as inputs to the GNN. (b) GRAS4T designs a graph contrastive learning framework, employing a GNN-based encoder, a GNN-based decoder, and a subspace module. This method effectively extracts potential representation via contrastive learning loss and reconstruction loss. (c) The potential embeddings learned through GRAS4T are essential for domain identification, trajectory inference, and are informative for downstream differential expression analysis.

### Data preprocessing and graph construction

The spatial transcriptomics data are preprocessed before converting them to graph structure data for the GRAS4T (Fig. 1a). Specifically, the top 3,000 variable genes are selected based on the raw gene expression profiles^14^. It should be noted that in the STARmap dataset, with less than 3,000 genes, no gene filtering is applied. For the MERFISH dataset, blank genes and Fos genes are removed. Additionally, cells labeled with ‘Ambiguous’ are also filtered out by following^27^. The selected gene expressions of top genes are then normalized based on the library and subsequently log-transformed. We denoted the preprocessed gene expression data as **X** *∈* ℝ^*d×n*^. The spatial location information and the gene expression are used to construct the adjacency matrix between the spots **A** *∈* ℝ^*n×n*^ where **A**_*ij*_≠ 0 if a linkage exists between spot i and spot j, otherwise **A**_*ij*_ = 0. By taking both spatial information and gene expression into consideration, GRAS4T selects the k nearest neighbors to construct the adjacency matrices **A**_*S*_, **A**_*E*_, respectively. These two matrices are merged using a balancing parameter α, expressed as **A** = **A**_*S*_ + α**A**_*E*_. Accordingly, the input graph 𝒢 can be described by the gene expression matrix **X** and the adjacency matrix **A**.

### Domain knowledge-aware graph augmentation module

Graph augmentation is one of the major components of GRAS4T (Fig. 1a). It typically involves attribute masking and edge perturbation^28^, both of which are random augmentations on a global scale (Supplementary Section 2.1). Inspired by^16, 29^, a domain knowledge-aware graph augmentation strategy is designed. Specifically, GRAS4T utilizes H&S image information to randomly perturb the edges from input graph 𝒢, thereby preserving information relevant to domain segmentation.

To obtain the morphology feature of the tissue, GRAS4T tiles the H&S image into the patches according to the spot distribution and uses Convolutional Neural Network to extract the latent representation **X**_*I*_. GRAS4T employs *k* nearest neighbor to construct the nearest neighbor graph **A**_*I*_, which then guides the augmentation of the nearest neighbor graph **A**. The graph augmentation function τ(·) is applied to graph 𝒢 to obtain the augmentation view. Here, the biological prior-based edge perturbation 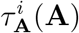 is defined as

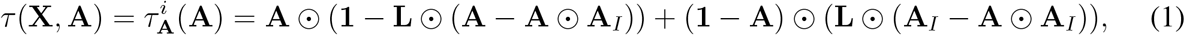

where ⊙ is the Hadamard product (element-wise product). **L** *∈* ℝ^*n×n*^ is a random perturbation location matrix, **L**_*ij*_ = **L**_*ji*_ = 1, if the connection between spot i and spot j is perturbed, otherwise **L**_*ij*_ = **L**_*ji*_ = 0. The two terms in 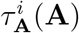 represent edge drop and addition in **A**, respectively. The first item represents a randomly disconnected link between spots with dissimilar tissue structures but similar spatial locations or gene expression, and the second item represents a randomly connected link between spots with similar tissue structures but not adjacent spatial locations or gene expression.

Accordingly, for the positive views, random graph augmentation methods proposed by You et al.^28^ and the prior-based graph augmentation mentioned above are used, i.e., the positive augmentation graphs 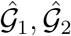obtained by graph augmentation in given graph 𝒢, where 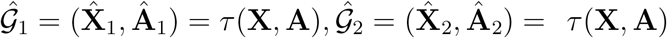. For the negative view 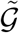, attribute shuffling is used to generate it (Supplementary section 2.1).

### Domain feature extraction module

The domain feature extraction module is another important component of GRAS4T (Fig. 1b), comprising an encoder, a symmetrical decoder, and a subspace model. Accordingly, it is trained by minimizing local-global contrastive loss, local-subspace contrastive loss, and reconstruction loss.

#### Graph representation learning

Graph contrastive learning^30–32^ is currently one of the most advanced methods in unsupervised representation learning, alleviating the heavy reliance on label information. The graph contrastive learning framework learns representation by maximizing the similarity between positive views while minimizing the similarity to negative views^16^. Deep Graph InfoMax (DGI)^30^, a well-known model in this field, discriminates nodes generated with correct topology and those generated with corrupted topology^33^ to make node representation contain as much global information as possible (Supplementary section 2.1). GRAS4T utilizes DGI as its foundational framework, integrating task-specific information and additional decoder structure.

Graph Convolutional Network (GCN) is used as the encoder of the GRAS4T. The single-layer encoder can be expressed as

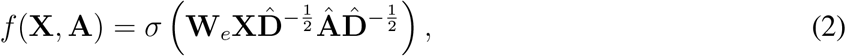

where **Â** = **A** + **I** is the adjacency matrix with inserted self-loops **I** and 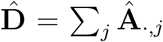 is the diagonal degree matrix. σ(·) is the nonlinear activation function (here we used Parametric Rectified Linear Unit^34^) and is applied column-wisely. **W**_*e*_ *∈* ℝ^*d′×d*^ (d′ is the dimension of hidden feature) is a learnable weight matrix. The representation **H** of the graph can be obtained by the GCN-based encoder.

Contrary to DGI which learns spot representation by contrasting local and global information in the original view, task-aware embeddings are learned by contrasting the local and global information of augmentation views. Thus, the local-global contrastive loss can be formulated as

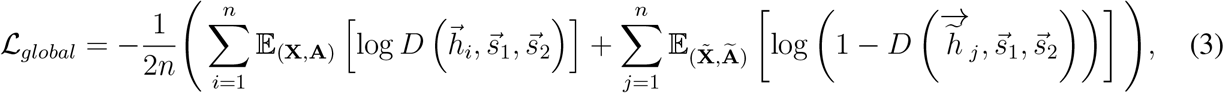

where 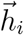 and 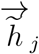 are the *i*-th spot of **H** and j-th spot of 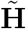, respectively. 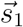 and 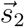 denote the global representation of the augmented graphs **Ĥ** _1_ and **Ĥ** _2_ accordingly. That it, 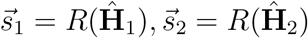, which represents the high-level summaries of augmented graph 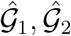. Here, R(·) is a readout function and D(·, ·, ·) is a discriminator, which outputs probability scores (defined in Supplementary section 2.1).

The structure of the encoder-decoder has the ability to preserve local spatial domain information^11, 13, 14^. In addition, the graph autoencoder provides local information while subspace-enhanced graph contrastive learning provides global and contextual information, which helps the GRAS4T model adapt to different downstream tasks. Here, we considered a GCN-based decoder symmetric with an encoder to reconstruct data **X**. The equation for the single-layer GCN-based decoder is

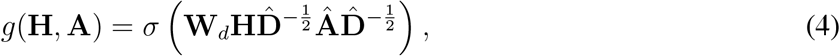

where **W** *∈* ℝ^*d×d′*^ is also a learnable matrix. The latent representation **H** is input to the decoder, which produces the reconstructed data **X**′.

To obtain an effective latent representation, the reconstruction loss between **X** and **X**′ is measured using mean-square error. The reconstruction loss is defined as

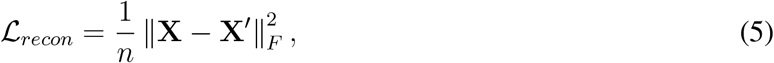

where ∥·∥_*F*_ denotes the Frobenius norm, i.e., ∥**X**∥_*F*_ =(∑_*ij*_ |x_*ij*_|^2^)^1*/*2^.

#### Subspace analysis

Subspace analysis^35–38^ is an important unsupervised learning method, which has achieved great success in bioinformatics, such as cell-type clustering^39, 40^ in single-cell transcriptomics and spatial domain identification in spatial transcriptomics^41^. As a state-of-the-art clustering method, subspace analysis is based on the self-expression assumption, which assumes that each data point can be represented by a linear combination of other data points in the same subspace^35^. More specifically, the key in subspace analysis is solving the following self-expression optimization problem

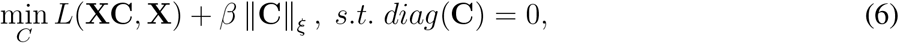

where **X** is the dataset and β is the balance parameter. **C** *∈* ℝ^*n×n*^ is the self-expression coefficient matrix and the *diag*(**C**) = 0 enforces a strict diagonal constraint on **C**. The first term of the objective function reconstructs each sample from all other samples, which describes the difference between the input data and the represented data. The second term introduces a regularisation of the coefficient matrix and different norms ξ are selected to obtain a self-expressive matrix with specific reconstruction properties. The subspace method relies on the assumption that data distribution is the union of subspace, however, ST data is complex and the relationships between spots are difficult to express linearly. To overcome the limitations above, the kernel-based method uses implicit nonlinear mappings to transfer the data into a space where the linearity assumption is more likely to be satisfied^42^.

Based on subspace theory, we used the self-expression matrix **C** to reconstruct the output of GCN-based encoder **Ĥ** _1_ and **Ĥ** _2_. More specifically, the self-expression matrix is obtained by solving the following optimization problem

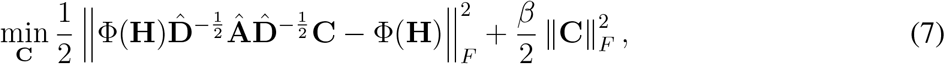

where Φ: ℝ^*m*^ *→ ℋ* is a mapping from the input space to the reproducing kernel Hilbert space *H*. The optimization problem mentioned above has a closed-form solution that is computationally efficient compared to some iterative numerical methods such as the alternating direction method of multipliers^43^. Meanwhile, to further enhance the acquisition of more closely connected co-domain nearest neighbors and remove residual noise from the ST data, a post-processing strategy was designed. To elaborate, we designed a filtering matrix **M** such that **M**_*i,j*_ = 1 if **C**_*i,j*_ is among the top k values in the j-th column of matrix **C** and **M**_*i,j*_ = 0 otherwise. We then multiplied this filtering matrix with the self-expression matrix to derive the subspace co-domain neighbor matrix **C**^*\**^. More details of this subspace module can be found in the Supplementary section 2.2.

Once the self-expression coefficient matrix **C** is obtained, a reconstructed representation containing local spatial information can be computed, i.e., **Z**_1_ = **Ĥ** _**1**_**C, Z**_2_ = **Ĥ** _**2**_**C** (the process can be considered as linearization^44^). Given that spots within the same subspace tend to belong to the same domain, the reconstructed representation encapsulates local information specific to that domain. We compared the reconstructed representation **Z**_1_ and **Z**_2_ containing neighbor information with the representation **H** and 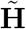, respectively. These comparisons led to the following local-subspace contrastive loss that can be defined by

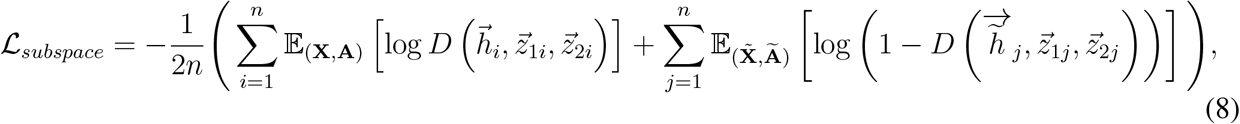

where 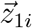 is the *i*-th spot of reconstructed representation **Z**_1_, which denote the contextual information of spot i from augmented graph 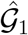. Similarly, the meaning of 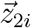 can be obtained.

#### Overall loss function

Combining Eq.(3), Eq.(5), and Eq.(8), GRAS4T optimized the following loss function

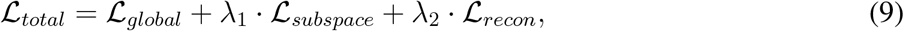

where λ_1_ and λ_2_ are the trade-off parameter that controls the balance between the effect of different modules.

### Downstream tasks

#### domain identification

GRAS4T employed two classical clustering methods to cluster low-dimensional representation, namely the mclust algorithm from the mclust R package^5^ and the Louvain algorithm from the scanpy package^45^. mclust and Louvain show advantages in datasets with different distributions. With the development of automated parameter selection tools, both clustering methods can be used in both annotated and unannotated datasets.

#### trajectory inference

The trajectory inference analysis was performed based on the clustering results of GRAS4T as grouping in the learned low-dimensional representations, using the Partition-based Graph Abstraction (PAGA)^46^ method from the scanpy package.

#### differential expression analysis

To identify domain-specific marker genes from the differential gene expression analysis across different domains, the ‘*correlation_matrix*’ and ‘*rank_genes_groups_dotplot*’ methods in the scanpy package were used. When using the ‘*rank_genes_groups_dotplot*’ method, ‘min_logfoldchange’ is set to 2.

### Parameter setting

GRAS4T is implemented via PyTorch 1.8.0. In the experiments, the balance parameter α between the two nearest neighbor graphs was adjusted according to the downstream clustering task. This parameter is set to 0.5 by default, and to 0.9 in the MERFISH dataset for cell type identification. The default graph augmentation methods for GRAS4T are ‘mask’ for nodes and ‘HS_image’ for edges. When the ST data does not contain an image, a random edge augmentation method ‘edge’ was used. In our study, the encoder was configured as a single-layer GCN by default, with a low-dimensional embedding dimension of 64. The decoder was consistent with the encoder structure. Although GRAS4T allows adjustments to the number of network layers and the dimensions of hidden layers, it is recommended to limit the GCN layers to a maximum of 3. Going beyond this recommendation often results in an over-smoothing effect, leading to poor expressivity^47^. Our model was optimized by the Adam optimizer^48^ with a learning rate of 0.001. GRAS4T alternated the model training and subspace optimization every 100 iterations until the maximum number of iterations was reached (500 iterations by default). For the subspace module, we used the default parameters provided by Cai et al.^37^. The ratio of global loss, subspace loss, and reconstruction loss in the final loss function is 1:1:1 by default. All experiments were implemented using NVIDIA GeForce RTX 3070Ti GPU, Intel® Core (TM) i7-12700K CPU @ 3.60 GHz, and 64 GB memory.

### Evaluation metrics

We used two evaluation metrics, Adjusted Rand Index^49^ (ARI) and Normalized Mutual Information^50^ (NMI) to quantify the similarity between cluster labels and manual annotations. These two evaluation metrics measure the clustering performance by measuring the similarity between the clustering results 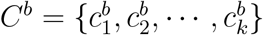 and the reference results 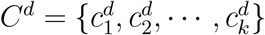. The cross-tabulation of C^*b*^ and C^*d*^ is shown in Table 1.

**Table 1.** The cross-tabulation of C^*b*^ and C^*d*^.

| | $c_1$ | $c_2$ | … | $c_k$ | sum |
| --- | --- | --- | --- | --- | --- |
| $c_1$ | $n_{11}$ | $n_{12}$ | … | $n_{1k}$ | $n_{1\cdot}$ |
| $c_2$ | $n_{21}$ | $n_{22}$ | … | $n_{2k}$ | $n_{2\cdot}$ |
| ⋮ | ⋮ | ⋮ | $\ddots$ | ⋮ | ⋮ |
| $c_k$ | $n_{k1}$ | $n_{k2}$ | … | $n_{kk}$ | $n_{k\cdot}$ |
| sum | $n_{\cdot 1}$ | $n_{\cdot 2}$ | … | $n_{\cdot k}$ | $n$ |

The ARI is a refined version of the Rand Index (RI). The RI treats the clustering results as a series of pairwise decisions and measures the clustering results based on the percentage of decisions that are correct. However, the RI cannot ensure that the score values for clustering results from randomized divisions are consistently near zero, this limitation led to the development of the ARI. The ARI score is calculated as

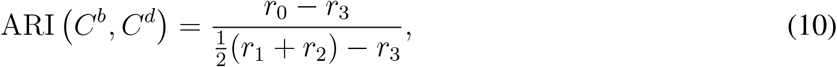

where

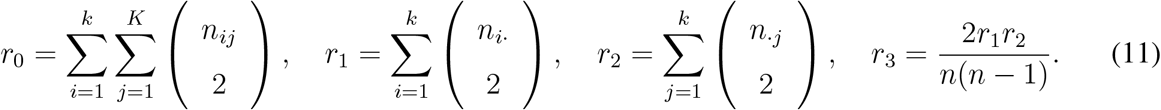

The NMI calculates the normalized similarity between two labels of the same data as follows

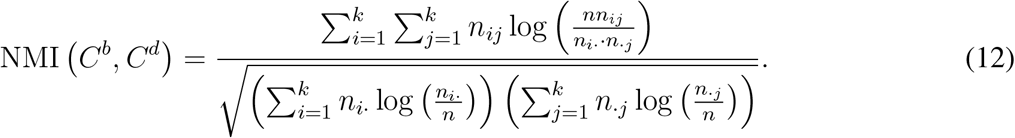

## Results

### Comparison GRAS4T with other methods on benchmark datasets

The effectiveness of the GRAS4T was first demonstrated by analyzing the DLPFC^19^ dataset and by comparing GRAS4T with other spatial domain identification methods including DeepST, STAGATE, SpaceFlow, conST, and CCST. GRAS4T demonstrated robust clustering capabilities and achieved the highest median ARI score across all 12 samples (Fig. 2a and Supplementary Figure S1), followed by STAGATE and DeepST, both based on graph autoencoder; SpaceFlow, conST, and CCST, utilizing DGI, showed smaller median ARI scores. Specifically, the median ARI score of GRAS4T surpasses that of STAGATE by 2.17% and outperforms CCST by 9.37%. As an example, in the DLPFC section 151672 (Fig. 2b), GRAS4T reported the best clustering accuracy (ARI=0.7045) while other models achieved ARI value ranged from 0.47 to 0.5926 (Fig. 2c). It could identify domains consistent with the manually annotated structure; whereas, CCST failed to separate the WM and Layer6, and DeepST, STAGATE, SpaceFlow, and conST exhibit challenges in accurately identifying Layer3 and Layer4 (Fig. 2c). When evaluating the performance on the DLPFC dataset with NMI, we observed that the clustering results of GRAS4T retained strong competitiveness (Supplementary Figures S1 and S5). In addition, the removal of the subspace module leads to 2.36% decrease in the median ARI score of GRAS4T (Supplementary Figure S2a). This observation demonstrates the significant role of the subspace module in enhancing clustering performance by effectively capturing local information and positively influencing the overall representation learning model. Concurrently, adopting random edge perturbation for the edge augmentation portion led to 0.83% decline in the median ARI score of GRAS4T (Supplementary Figure S2b).

**Figure 2.**
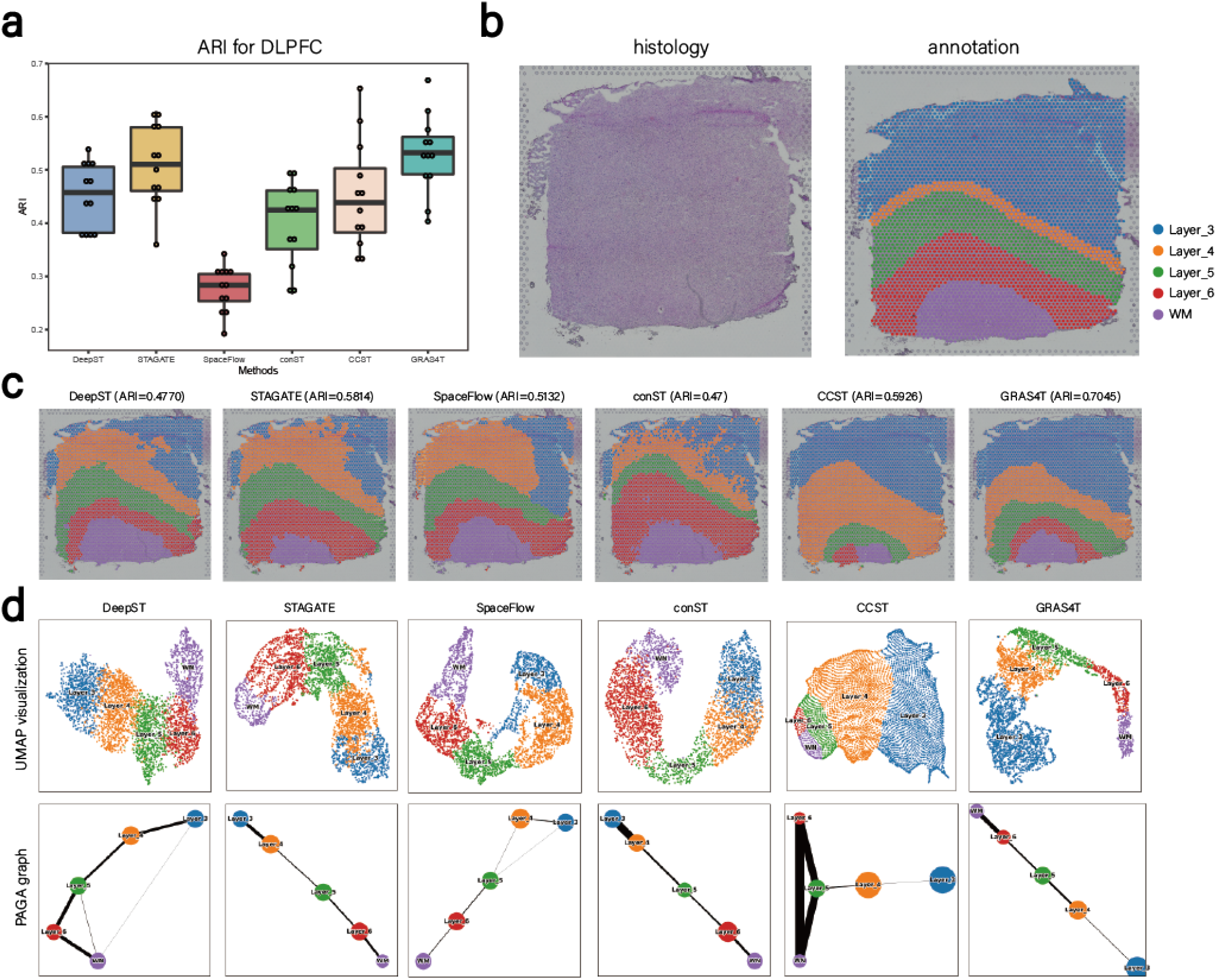
GRAS4T enhanced the precision of identifying layer structures within the DLPFC dataset. (a) Boxplot of clustering accuracy in all sections of the DLPFC dataset in terms of ARI values for six methods. (b) The H&E image and manual annotation. (c) The spatial clusters in six methods for slice 151672. (d) UMAP visualizations and PAGA graph in six methods for slice 151672.

Notably, the visualization of the low-dimensional embedding of GRAS4T using Uniform Manifold Approximation and Projection (UMAP)^51^ demonstrated spatial trajectories that were consistent with the developmental scenario (Fig. 2d and Supplementary Figure S3). Using slice 151672 as an example, the UMAP of the embeddings successfully illustrated the developmental trajectory from WM to Layer 6. In contrast, CCST was unable to clearly depict the developmental hierarchy from WM to Layer 6. The well-revealed trajectory by the UMAP plot of GRAS4T was further confirmed through the application of PAGA, which delineated the developmental trajectories across the various layers (Fig. 2d and Supplementary Figure S3). The results elucidated that among the algorithms evaluated, only GRAS4T, STAGATE, and conST, produced PAGA graphs that unambiguously captured trajectories aligning with the anticipated developmental trajectories.

The efficacy of GRAS4T was further evaluated using the human HER2-positive breast tumor (HER2+) dataset^21^, which used different spatial technologies (spatial transcriptomics) than the DLPFC dataset. We noticed a significant performance advantage of GRAS4T over all the other benchmark methods across the eight slices (A-H) in the HER2+ dataset (Fig. 3a and Supplementary Figures S4). Still, the median ARI score of GRAS4T is 0.3468, while that of all other methods fall below 0.3. Notably, GRAS4T’s median ARI surpasses that of the second-ranked CCST method by 7.05%. STAGATE closely follows, with SpaceFlow and conST exhibiting comparable performances. DeepST recorded the lowest median ARI score among the methods evaluated. For the A1 section, GRAS4T’s identification aligned most closely with manual annotation shapes, achieving the highest ARI score (ARI=0.6349) as shown in Fig. 3b and c. Importantly, we accurately identified both invasive cancer and *in situ* cancer. In comparison, STAGATE mixed spots from different domains, whereas CCST, despite demonstrating more distinct cluster delineation, it misidentified certain areas of invasive cancer as connective tissue. When employing NMI to assess the performance on the HER2+ dataset, GRAS4T demonstrated an advantage, indicating its efficacy in achieving robust spatial domain identification, even within the context of class imbalance in the ST dataset (Supplementary Figures S4 and S5). We further tested the performance of six methods on a human breast cancer dataset from the 10X Visium platform. Compared with other spatial algorithms, GRAS4T showed a more comprehensive identification of the ductal carcinoma (DCIS) region, the lobular carcinoma (LCIS) region, and the invasive ductal carcinoma (IDC) region. Notably, GRAS4T achieved 0.83% higher median ARI than CCST, the second-ranked method (Supplementary Figure S6).

**Figure 3.**
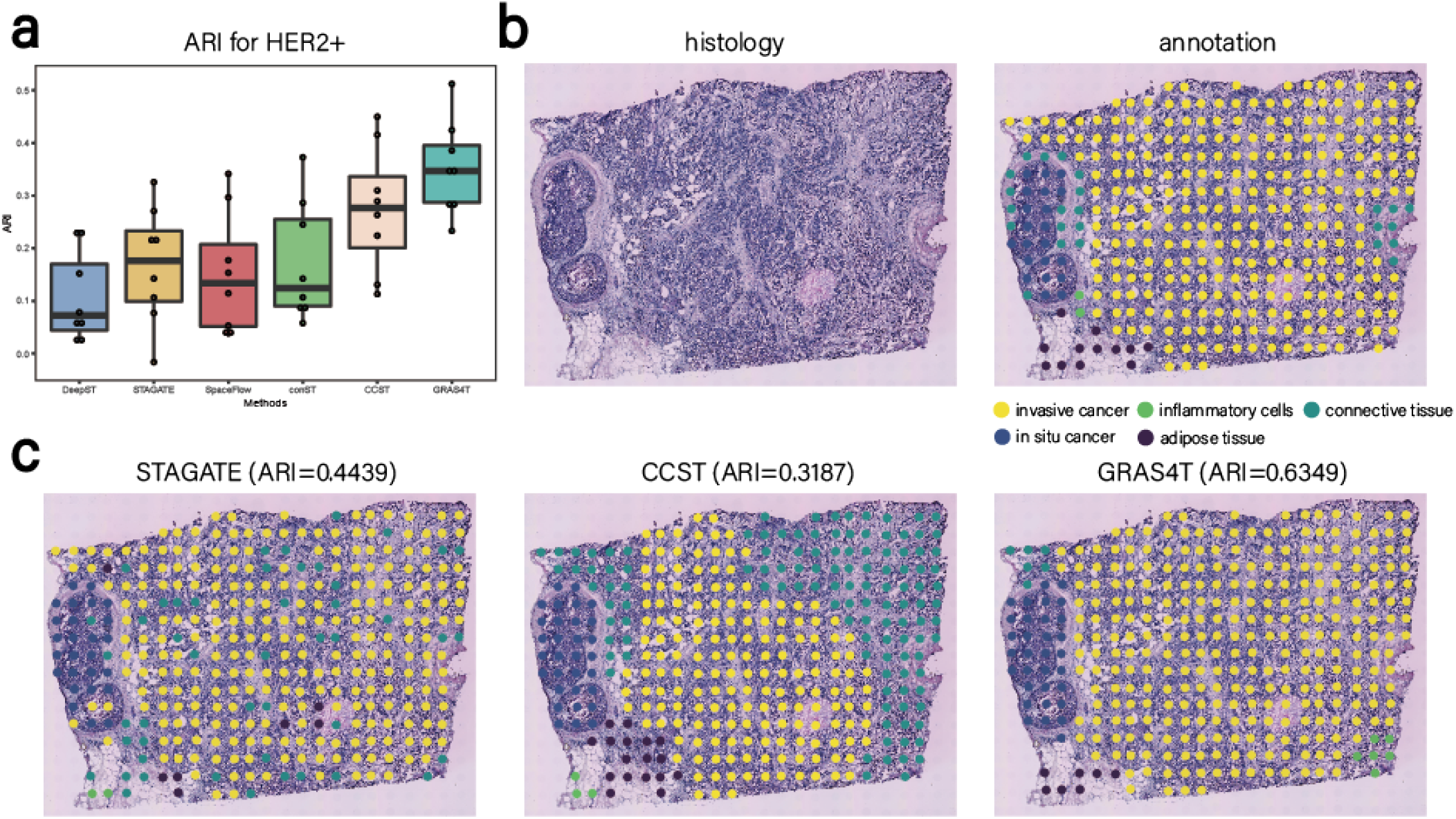
GRAS4T accurately discovered spatial domains in the HER2+ dataset. (a) Boxplot of clustering accuracy in all sections of the HER2+ dataset in terms of ARI values for all methods. (b) The H&E image and manual annotation for the A1 section. (c) Spatial domains of HER2+ data in (b) detected by STAGATE, CCST, and GRAS4T.

### GRAS4T enhances domain separation with its block diagonal property

The mouse visual cortex dataset was used to demonstrate the spatial domain separation capability of GRAS4T. GRAS4T accurately identified the Hippocampus (HPC), Corpus Callosum (CC), and various layers of the cerebral cortex (L1, L2/3, L4, L5, and L6) in the mouse visual cortex. Compared to STAGATE and CCST, the domain regions identified by GRAS4T are more contiguous and highly consistent with manual labeling (Fig. 4a). It is evidenced through the ARI pirate plots of the six methods that GRAS4T has the highest median score. STAGTE, CCST, and SpaceFlow have similar identification performance, while DeepST and conST have poor identification results (Fig. 4b). The UMAP visualization of the embedding results demonstrated the trajectory from HPC, which is located in the deep part of the brain, to CC, and subsequently from L6 to L1, the most superficial layer of the cerebral cortex. This pattern of change accurately describes the differentiation process in the mouse brain (Fig. 4c). To gain insight into the gene expression profiles across distinct brain regions, we performed differentially expressed gene analysis for each domain cells against the remaining cells. The top three domain specific genes per domain were illustrated in the dotplot (Fig. 4d). Our results revealed that the *3110035E14Rik* gene exhibits elevated expression in the L6 layer, while the *Cplx1* gene demonstrates high expression in the L5 layer. These results align with previous reports^23^, bolstering the validity of our observations and providing additional support to the existing knowledge (Fig. 4d).

**Figure 4.**
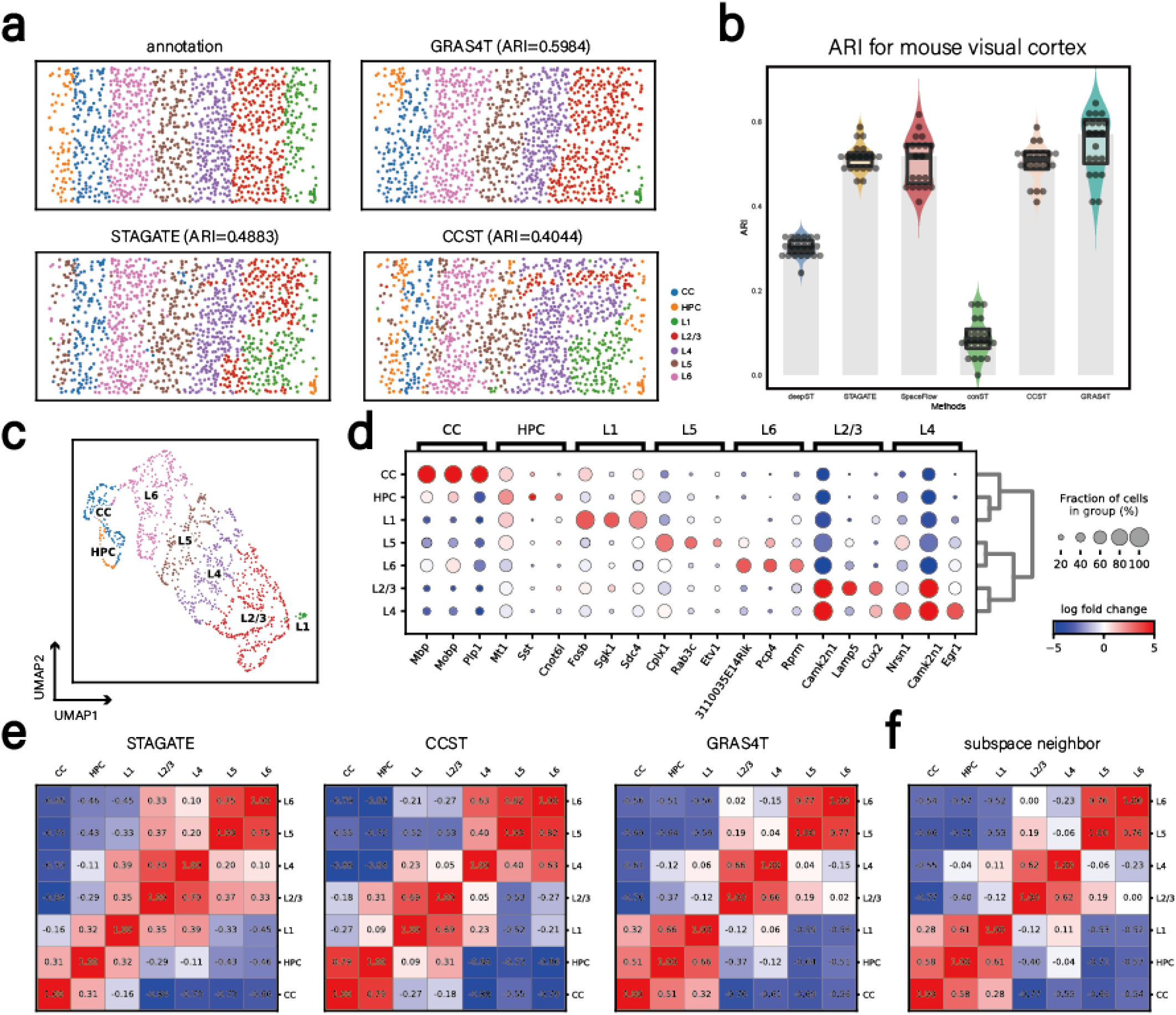
GRAS4T improved spatial domain identification results in the mouse visual cortex dataset and enhanced the separation between different spatial domains. (a) The manual annotation on mouse visual cortex and spatial domains were detected by GRAS4T, STAGATE, and CCST. (b) Comparison of different methods by ARI pirate graph. (c) UMAP visualization using representations generated by GRAS4T. (d) Dotplot of the top 3 domain-specific genes for each domain. The size of the dots represents the proportion of cells expressing the gene. The color represents the average expression of the gene in the region where red reflects high expression and blue means low expression. (e) Heatmap of inter-domain Pearson correlation coefficients for STAGATE, CCST, and GRAS4T. (f) Heatmap of inter-domain Pearson correlation coefficients for subspace nearest neighbors.

In our pursuit of unraveling the inter-domain heterogeneity of gene expression, we calculated the inter-domain Pearson correlation coefficients for each of the three methods. Compared to STAGATE, the heatmaps of GRAS4T and CCST show more dissimilarity in the non-diagonal sections. Taking GRAS4T, which has the highest ARI, as a benchmark, we found significant heterogeneity between CC, HPC, and L1 and other domains (Fig. 4e). To observe the nearest neighbor coverage of individual spots, we assigned the most frequent class in the subspace nearest neighbors of each spot as its class and generated the correlation heatmap. The results indicate that the correlation coefficient heatmap of subspace nearest neighbors exhibits a more block-diagonal pattern, suggesting that the spatial domains optimized by these neighbors likewise show stronger spatial heterogeneity (Fig. 4f). It is worth mentioning that our method consistently exhibited superior spatial heterogeneity in section 151672 of DLPFC and A1 of HER2+, surpassing that of STAGATE and CCST (Supplementary Figure S7). In the following, we will show further advantages of GRAS4T in the task of deciphering spatial domains with more datasets. This performance was benchmarked against STAGATE and CCST, which are exemplary methods in the field of spatial domain identification.

### GRAS4T captures co-domain neighbor to reveal finer-grained tissue structures

The GRAS4T’s capacity was assessed to select co-domain neighborhoods in complex biological tissues, particularly, on the coronal mouse brain dataset (Fig. 5a). By utilizing the subspace module to capture localization information, we found that GRAS4T was able to identify smaller spatial domains in the hippocampus, such as CA1 (domain 6), CA3 (domain 19), and dentate gyrus (domain 13). By incorporating H&S images for graph augmentation, GRAS4T was able to identify regions that better matched the boundary information in morphological images. Although STAGATE could also describe small regions in the hippocampal part, it was not as effective in segmenting the boundaries of morphological images. Furthermore, CCST was unable to depict small spatial domains despite being able to smooth domain boundaries. It’s worth noting that GRAS4T without subspace selection of nearest neighbors failed to identify domain 13 and domain 19, which were identified similarly to CCST (Fig. 5b).

**Figure 5.**
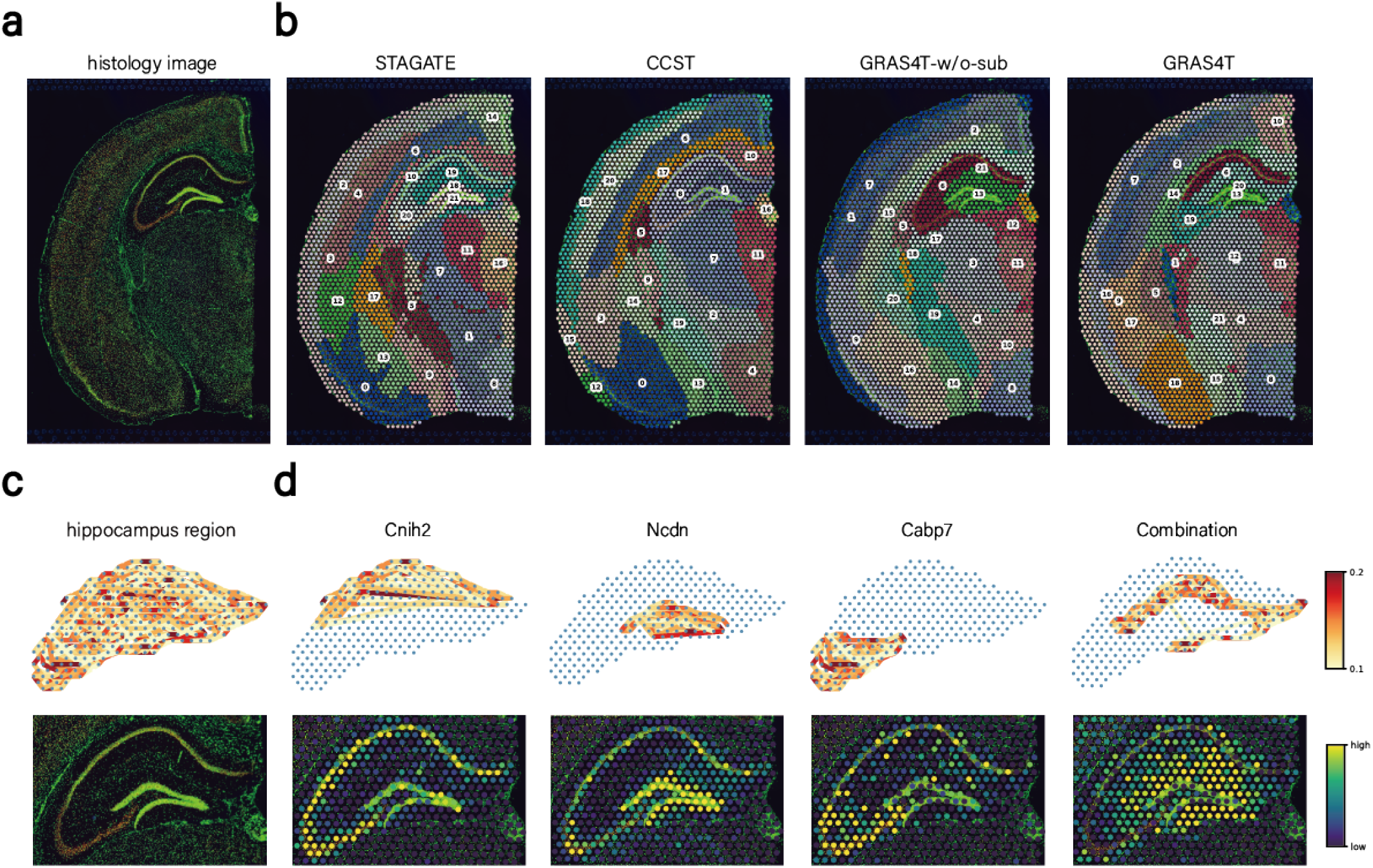
GRAS4T revealed refined spatial domains in the coronal section of an adult mouse brain. (a) The coronal mouse brain image stained with DAPI and Anti-NeuN. (b) Spatial domains generated by STAGATE, CCST, and GRAS4T without or with subspace module. (c) Visualization of the affinity matrix and H&S image for the hippocampal region. The nodes in the graph were reflected by the spots and the edges were determined by the values in the affinity matrix. (d) The visualization of spatial domains identified by GRAS4T for the corresponding marker genes.

In order to delve deeper into the potential of GRAS4T’s co-domain nearest neighbor selectivity to reveal finer tissue structures, we directed our investigation towards the hippocampal region and constructed a network heatmap utilizing spot coordinates and subspace neighbor information (Fig. 5c-d). Examining the network of non-convex domain 6 and domain 20, it was evidenced that GRAS4T selects spots that were not necessarily closely positioned physically but were located in the same domain as nearest neighbor points, which facilitates the discovery of complex structural regions. The spatial domains identified by GRAS4T were clearly supported by known gene markers. For example, *Ncdn* was strongly expressed on dentate gyrus^52^, while CA1 displays pronounced expression of *Cabp7*^53^. Notably, we identified a unique gene combination (Combination = *Ddn*+*Camk2a*+*Slc1a2*-*Ncdn*-*Cnih2*) that accurately characterizes domain 6. Collectively, these findings demonstrated the remarkable ability of the GRAS4T subspace module to reveal finer-grained tissue structures. We further tested the performance of STAGATE, CCST, and GRAS4T on the mouse brain anterior&posterior dataset. We observed that GRAS4T and STAGATE showed great agreement with reference (Supplementary Figure S8). In particular, GRAS4T can capture delicate structures like the dorsal and the ventral horn of the hippocampus region.

### GRAS4T discerns the anatomical regions of tissue from ST data with different spatial resolutions

GRAS4T exhibited a remarkable performance on identifying complex spatial domains on datasets from both the 10X Visium and STARmap platforms. Furthermore, GRAS4T could also be adapted to other ST datasets with different spatial resolutions. DAPI-stained image^54^ was employed to the mouse olfactory bulb dataset generated by Stereo-seq^25^ as a criterion for the spatial domain identification task (Fig. 6a). Although CCST can clearly segment the boundaries of each layer, it inappropriately divides the rostral migratory stream (RMS) into three domains and fails to delineate the inner layers, namely the mitral cell layer (MCL), internal plexiform layer (IPL), granule cell layer (GCL), and RMS. STAGATE was unable to clearly distinguish between the external plexiform layer (EPL) and glomerular layer (GL); the boundaries of layers were mixed. Compared with STAGATE and CCST, the GRAS4T detected olfactory nerve layer (ONL), GL, EPL, and MCL more distinctly, and also successfully identified the GCL (Fig. 6b). The performance of GRAS4T was further assessed in the mouse hypothalamus datasets generated by MERFISH^26^. Specifically, we performed a comparative analysis of STAGATE, CCST, and GRAS4T with reference cell types on 12 distinct slices of the mouse hypothalamus dataset. The experimental results showed that GRAS4T remains competitiveness in domain identification with discontinuous domains (Supplementary Figure S9). For example, GRAS4T accurately identified oligodendrocytes (OD Mature2 domain) in the +0.16 slice and successfully captured Ependymal domain in the −0.29 slide (Fig. 6c). Compared to STAGATE (ARI=0.0719 for +0.16, ARI=0.1167 for −0.29) and CCST (ARI=0.0494 for +0.16, ARI=0.0251 for −0.29), GRAS4T showed a high degree of consistency to the reference cell types (ARI=0.4641 for +0.16, ARI=0.4284 for −0.29).

**Figure 6.**
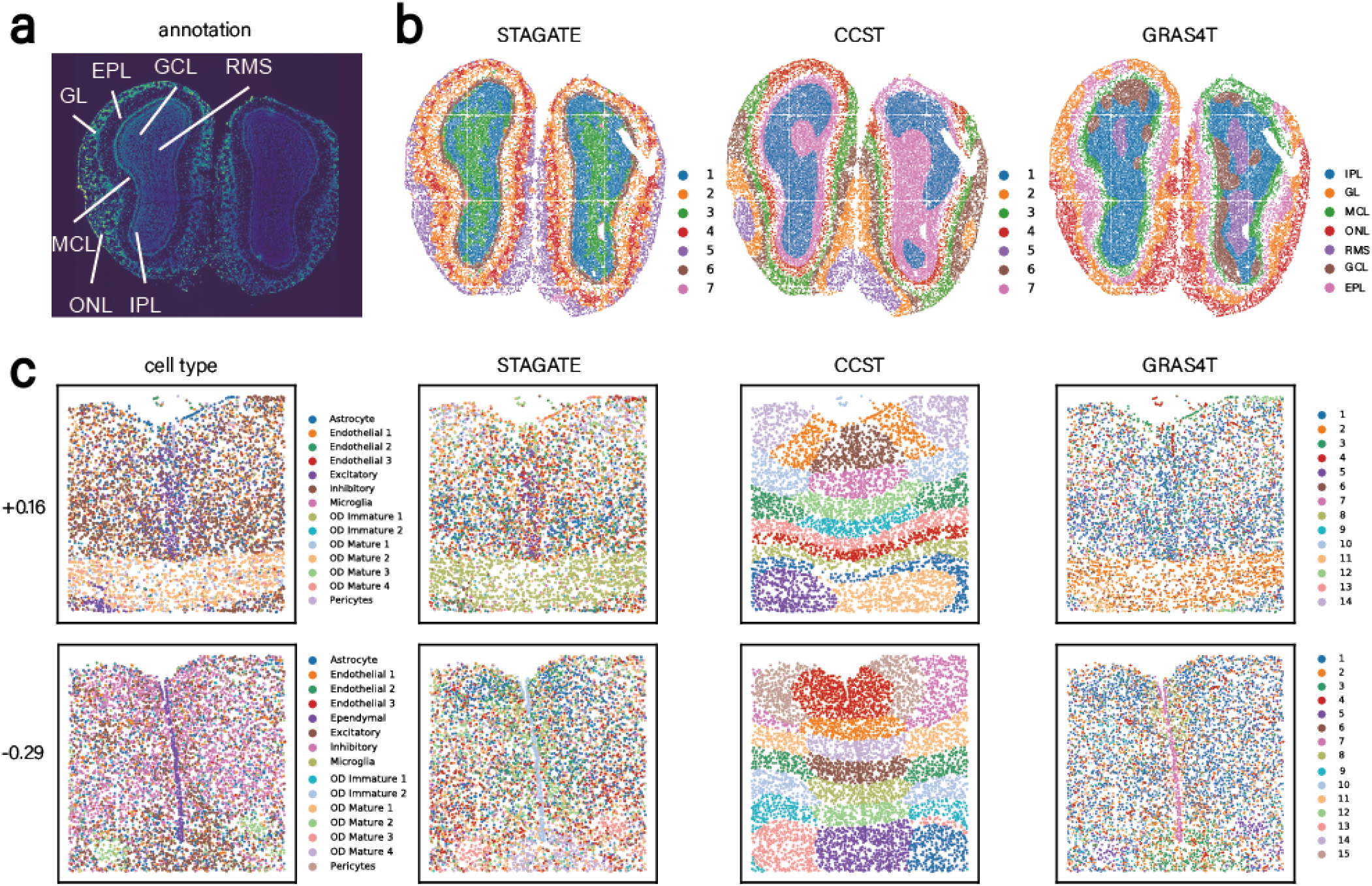
GRAS4T identifies spatial domains in ST datasets profiled by Stereo-seq and MERFISH platform. (a) Laminar tissue of mouse olfactory bulb annotated in the DAPI-stained image. (b) Spatial domains detected by STAGATE, CCST, and GRAS4T. (c) Cell type and spatial domain identification by STAGATE, CCST, and GRAS4T on the 2 slices of mouse hypothalamus datasets.

## Discussion

The emergence of spatial transcriptomics has provided a spatial depiction of biological tissue structures and cellular functions. Naturally, the rich relationships among spots can be effectively captured through graph modeling, and learning graph representation is crucial for understanding the heterogeneity and similarity among spots. In this study, we developed a graph contrastive learning framework GRAS4T to perform spatial clustering or identify domains in spatial transcriptomics. GRAS4T simultaneously considers the local environment of spots, spatial domain features, and global information of the tissue section, thereby enabling precise delineation of spatial domains and adapting to a variety of downstream tasks. Specifically, GRAS4T retains crucial information related to domain segmentation tasks in positive views through domain knowledge-aware graph augmentation. It also utilizes a subspace analysis module to adaptively capture the tissue microenvironment and spatial domain information. The domain knowledge-aware graph augmentation and subspace analysis strategy results in the effective separation of inter-domain regions and the precise capture of co-domain neighbors. We evaluated GRAS4T across a broad range of spatial transcriptomics datasets, encompassing different platforms and tissue types. The findings demonstrated the superiority of GRAS4T in identifying spatial domains.

One essential component of GRAS4T is the domain knowledge-aware graph augmentation in graph contrastive learning. Currently, the spots relationships in the image (or morphology features) are employed to perform augmentation for the positive views where interactions and relationships between cells in spatially proximate neighborhoods are emphasized. For those ST datasets without image data associated, random graph augmentations are adopted to show their effectiveness in augmenting positive views. We noted that cells within the inferred subspace modules generally show similar gene expression profile or furthermore potentially similar cell states or phenotypic characteristics, thus would be a good potential for positive view augmentation. Furthermore, interactions between cells along spatial gradients or domain boundaries might also be a potential angle for graph augmentation. Some other studies also suggest to seek for cells shared similar biological characterizations or annotations (domains) or functional modules from the same datasets or from replicates^55, 56^. In addition, when the spatial datasets have a temporal series, creating positive views which capture similar patterns or dynamic changes over times might be another effective method for graph augmentation. These merit further investigations for graph augmentation and also for multiple slices data integration.

GRAS4T incorporates a subspace module with a graph contrastive learning framework, leveraging the strengths of both machine learning and deep learning approaches. Indeed, the combination of machine learning and deep learning is an interesting topic in the AI field^57–59^. A representation learning framework is required to produce a well-structured data representation that is suitable for subspace analysis (i.e., the representations in the latent features space approximately lie in a union of subspaces)^60^. Concurrently, the representation with subspace structure can enhance the interpretability of deep learning models and mitigate the risk of model collapse^61^. Here, we use the self-expression matrix learned from the subspace to reconstruct the low-dimensional representations of the positive views through a one-step linearization^44^ and to obtain the representations of the microenvironments that contain the same domain tissue. Meanwhile, we provide a post-processing tool that could mask the self-expression matrix based on the magnitude of self-expression values^62^. This procedure effectively severs links between target points and non-domain counterparts, efficiently eliminating noise. Notably, this post-processing method can be characterized as an avenue for capturing the core cell set, facilitating the alignment and integration of multiple ST slices.

While our comprehensive comparison experiments show encouraging results, there is considerable potential for further development and extension of the GRAS4T. Besides the potential extension to multiple ST datasets integration, explorations of various regularization items for the subspace models are important for achieving promising performance. Different types of regularization confer distinct properties to the self-expression matrix. The selection of regularization would be contingent on the specific dataset and task as inappropriate regular terms could significantly impair the quality of reconstructed representations. Hence, considering the context of complex ST data and the diversity of downstream tasks, it is merit to further explore the impacts of different regularization methods such as elastic net regularization^63^, *l*_1_ regularization^35^, etc. Taken together, there exists substantial potentials to investigate the integration of subspace module and deep learning architectures as well as various regulation methods. This will eventually enhance our ability to more effectively tackle relevant biological questions such as trajectory inference and cell-cell communications.

In conclusion, this study introduces GRAS4T, an innovative graph contrastive learning framework designed to effectively identify spatial domains in spatial transcriptomics. GRAS4T integrates local spot information, contextual domain information, and global tissue information, enhancing its effectiveness and versatility in downstream applications. Central to GRAS4T’s efficacy is the domain knowledge-aware graph augmentation and the subspace module, both of which contribute to the effective separation of distinct spatial domains and the precise identification of co-domain neighbors. Through a detailed discussion about GRAS4T and a wide range of experiments on different ST data, we demonstrate that GRAS4T is an ideal framework with extensibility. In future works, we will develop this framework further based on the ST background to make it more versatile and interpretable.

## Data availability

The ST datasets supporting the findings of this study are all publicly available. (1) The DLPFC dataset is available at http://research.libd.org/spatialLIBD/. (2) The HER2+ dataset generated by spatial transcriptomics platform is accessed at https://github.com/almaan/her2st. (3) The mouse visual cortex dataset generated by STARmap is available at https://www.dropbox.com/sh/f7ebheru1lbz91s/AADm6D54GSEFXB1feRy6OSASa/visual_1020/20180505_BY3_1kgenes?dl=0&subfolder_nav_tracking=1. (4) The adult mouse brain dataset is accessed at https://www.10xgenomics.com/resources/datasets. (5) The Stereo-seq mouse olfactory bulb dataset is available at https://github.com/JinmiaoChenLab/SEDR_analyses/. (6) The MERFISH dataset is accessed at https://datadryad.org/stash/dataset/doi:10.5061/dryad.8t8s248. (7) The human breast cancer dataset is available at https://www.10xgenomics.com/resources/datasets. (8) The anterior and posterior sections of the mouse brain are accessed at https://www.10xgenomics.com/resources/datasets and the Allen Brain Atlas reference is available at https://mouse.brain-map.org/static/atlas.

## Code availability

All source codes used in our experiments have been deposited at https://github.com/Lab-Xu/GRAS4T.

