## Supplementary for "Spatial domains identification in spatial transcriptomics by domain knowledge-aware and subspace-enhanced graph contrastive learning"

The supplementary material contains experimental settings, details of GRAS4T, and additional experimental results.

### 1 Experimental Settings

In this section, we described the baseline datasets,compared methods, and evaluation metrics in this paper.

#### 1.1 datasets

We presented eight sets (38 sections in total) of challenging spatial transcriptomic (ST) datasets that were utilized in this paper. These datasets covered different platforms, tissues, sizes, and number of clusters, as detailed in Table S1.

Table S1: Detailed description of the datasets in the paper.

| Plarform | Tissue | Section | Spot | Gene | Cluster | Reference |
|---|---|---|---|---|---|---|
| 10X Visium | human dorsolateral prefrontal cortex | 151507 | 4226 | 33538 | 7 | [1] |
| | | 151508 | 4384 | 33538 | 7 | |
| | | 151509 | 4789 | 33538 | 7 | |
| | | 151510 | 4634 | 33538 | 7 | |
| | | 151669 | 3661 | 33538 | 5 | |
| | | 151670 | 3498 | 33538 | 5 | |
| | | 151671 | 4110 | 33538 | 5 | |
| | | 151672 | 4015 | 33538 | 5 | |
| | | 151673 | 3639 | 33538 | 7 | |
| | | 151674 | 3639 | 33538 | 7 | |
| | | 151675 | 3673 | 33538 | 7 | |
| | | 151676 | 3460 | 33538 | 7 | |
| | human breast cancer | - | 3798 | 36601 | 20 | - |
| | mouse brain | coronal | 2903 | 32285 | - | - |
| | | anterior | 2823 | 32285 | - | [2] |
| | | posterior | 3289 | 32285 | - | |
| spatial transcriptomics | human breast cancer | A1 | 341 | 15045 | 5 | [3] |
| | | B1 | 269 | 15109 | 4 | |
| | | C1 | 167 | 15557 | 3 | |
| | | D1 | 255 | 15661 | 3 | |
| | | E1 | 534 | 15701 | 3 | |
| | | F1 | 659 | 14861 | 3 | |
| | | G2 | 402 | 15258 | 6 | |
| | | H1 | 530 | 15029 | 6 | |
| MERFISH | mouse hippocampus | -0.29 | 5517 | 155 | 15 | [4] |
| | | -0.24 | 5543 | 155 | 15 | |
| | | -0.19 | 5803 | 155 | 15 | |
| | | -0.14 | 5926 | 155 | 15 | |
| | | -0.09 | 5557 | 155 | 15 | |
| | | -0.04 | 6154 | 155 | 14 | |
| | | +0.01 | 5338 | 155 | 15 | |
| | | +0.06 | 5343 | 155 | 15 | |
| | | +0.11 | 5070 | 155 | 15 | |
| | | +0.16 | 6067 | 155 | 14 | |
| | | +0.21 | 4787 | 155 | 15 | |
| | | +0.26 | 4832 | 155 | 15 | |
| Stereo-seq | mouse olfactory bulb | - | 19109 | 27106 | 7 | [5] |
| STARMAP | mouse visual cortex | - | 1207 | 1020 | 7 | [6] |

#### 1.2 compared methods

We compared GRAS4T with the following state-of-the-art spatial domain identification methods: DeepST, STAGATE, conST, SpaceFlow, and CCST. For algorithmic comparisons, we employed the default parameters provided by these methods in their publications and codebases. To ensure fairness, we engaged hyperparameter tuning to rectify evidently problematic outcomes in the identification results. The software packages for all the baseline methods utilized in this paper are listed in Table S2.

Table S2: Detailed description of the software packages of baseline methods.

| Method | Language | Access | Reference |
|---|---|---|---|
| DeepST | python | https://github.com/JiangBioLab/DeepST | [7] |
| STAGATE | python | https:// github.com/zhanglabtools/STAGATE | [8] |
| conST | python | https://github.com/ys-zong/conST | [9] |
| SpaceFlow | python | https://github.com/hongleir/SpaceFlow | [10] |
| CCST | python | https://github.com/xiaoyeye/CCST | [11] |

#### 1.3 evaluation metrics

We used two evaluation metrics, Adjusted Rand Index [12] (ARI) and Normalized Mutual Information [13] (NMI) to quantify the similarity between cluster labels and manual annotations. These two evaluation metrics measure the clustering performance by measuring the similarity between the clustering results $C^b = \{c_1^b, c_2^b, \cdots, c_k^b\}$ and the reference results $C^d = \{c_1^d, c_2^d, \cdots, c_k^d\}$. The cross-tabulation of $C^b$ and $C^d$ is shown in Table S3.

Table S3: The cross-tabulation of $C^b$ and $C^d$.

| | $c_1$ | $c_2$ | $\cdots$ | $c_k$ | sum |
|---|---|---|---|---|---|
| $c_1$ | $n_{11}$ | $n_{12}$ | $\cdots$ | $n_{1k}$ | $n_{1\cdot}$ |
| $c_2$ | $n_{21}$ | $n_{22}$ | $\cdots$ | $n_{2k}$ | $n_{2\cdot}$ |
| $\vdots$ | $\vdots$ | $\vdots$ | $\ddots$ | $\vdots$ | $\vdots$ |
| $c_k$ | $n_{k1}$ | $n_{k2}$ | $\cdots$ | $n_{kk}$ | $n_{k\cdot}$ |
| sum | $n_{\cdot 1}$ | $n_{\cdot 2}$ | $\cdots$ | $n_{\cdot k}$ | $n$ |

The ARI is a refined version of the Rand Index (RI). The RI treats the clustering results as a series of pairwise decisions and measures the clustering results based on the percentage of decisions that are correct. However, the RI cannot ensure that the score values for clustering results from randomized divisions are consistently near zero, this limitation led to the development of the ARI. The ARI score is calculated as

$$\text{ARI}\left(C^b, C^d\right) = \frac{r_0 - r_3}{\frac{1}{2}(r_1 + r_2) - r_3}, \tag{1}$$

where

$$r_0 = \sum_{i=1}^{k} \sum_{j=1}^{K} \binom{n_{ij}}{2}, \quad r_1 = \sum_{i=1}^{k} \binom{n_{i\cdot}}{2},$$
$$r_2 = \sum_{j=1}^{k} \binom{n_{\cdot j}}{2}, \quad r_3 = \frac{2r_1 r_2}{n(n-1)}. \tag{2}$$

The NMI calculates the normalized similarity between two labels of the same data as follows

$$\mathrm{NMI}\left(C^b, C^d\right) = \frac{\sum_{i=1}^{k}\sum_{j=1}^{k} n_{ij}\log\left(\frac{n n_{ij}}{n_{i\cdot}\cdot n_{\cdot j}}\right)}{\sqrt{\left(\sum_{i=1}^{k} n_{i\cdot}\log\left(\frac{n_{i\cdot}}{n}\right)\right)\left(\sum_{j=1}^{k} n_{\cdot j}\log\left(\frac{n_{\cdot j}}{n}\right)\right)}}. \tag{3}$$

#### 2 Detail of GRAS4T

##### 2.1 graph contrastive learning

The graph contrastive learning framework consists of three main components:

- The graph data augmentation module is responsible for generating different views of a given graph. Graph data augmentation is essential in graph contrastive learning. When appropriate augmentations are applied, the corresponding prior is instilled and the model learns representations useful for downstream tasks by maximizing the consistency between the graph and its augmentations.
- The GNN-based encoder is used for computing representations. By training on data using Graph Neural Network (GNN), hidden representations are obtained, which contain rich graph structure information.
- The contrastive learning target is utilized for training the model. Contrastive learning obtains representations by maximizing mutual information between instances with similar semantic information, various pretext tasks can be constructed to enrich the supervision signals from such information [14].

The Deep Graph InfoMax (DGI) [15] is one of the most famous graph contrastive learning models. Given the undirected attribute graph $\mathcal{G} = \{\mathcal{V}, \mathcal{E}, \mathbf{X}\}$, where $\mathcal{V} = \{v_1, v_2, \cdots, v_n\}$, $(|\mathcal{V}| = n)$ is the set of nodes and $\mathcal{E}(|\mathcal{E}| = m)$ is the set of edges representing the relationship between node $i$ and node $j$. The neighbors of the node $v_i$ are denoted as $\mathcal{N}(v_i) = \{v_j \in \mathcal{V} \mid e_{i,j} \in \mathcal{E}\}$. $\mathbf{A} \in \mathbb{R}^{n\times n}$ is the adjacency matrix of $\mathcal{G}$ such that $\mathbf{A}_{i,j} = 1$ if $e_{i,j} \in \mathcal{E}$ and $\mathbf{A}_{i,j} = 0$ otherwise. $\mathbf{X} = [\boldsymbol{x}_1, \boldsymbol{x}_2, \cdots, \boldsymbol{x}_n] \in \mathbb{R}^{d\times n}$ is the feature matrix, where $d$ is the dimension of the node.

The graph augmentation $\tau(\cdot)$ is applied in graph $\mathcal{G}$ to obtain the augmentation view. Many graph data augmentation methods have been developed, such as attribute (feature) masking, edge perturbation, and attribute shuffling [14, 15]. The DGI obtains a negative view through attribute shuffling, i.e. $(\tilde{\mathbf{X}}, \tilde{\mathbf{A}}) = \tau(\mathbf{X}, \mathbf{A})$. The attribute shuffling performs the column-wise shuffling on the feature matrix, we specify $\tau^s_{\mathbf{X}}(\mathbf{X})$ for attribute shuffling as

$$\tau(\mathbf{X}, \mathbf{A}) = \tau^s_{\mathbf{X}}(\mathbf{X}) = \mathbf{X}\left[:, idx\right], \tag{4}$$

where $idx$ is a randomly ordered list containing numbers from 1 to $n$. Attribute shuffling serves as an effective method for corrupting the graph structure by using column swapping.

DGI utilizes a one-layer Graph Convolutional Network (GCN) as the encoder of the model, the propagation of the GCN is depicted as

$$f(\mathbf{X}, \mathbf{A}) = \sigma\left(\mathbf{W}_e\mathbf{X}\hat{\mathbf{D}}^{-\frac{1}{2}}\hat{\mathbf{A}}\hat{\mathbf{D}}^{-\frac{1}{2}}\right), \tag{5}$$

where $\hat{\mathbf{A}} = \mathbf{A} + \mathbf{I}_N$ is the adjacency matrix with inserted self-loops and $\hat{\mathbf{D}} = \sum_j \hat{\mathbf{A}}_{\cdot,j}$ is the diagonal degree matrix. $\sigma(\cdot)$ denotes the nonlinear activation function such as the Parametric Rectified Linear Unit, applied column-wise. $\mathbf{W}_e \in \mathbb{R}^{d'\times d}$ ($d'$ is the hidden feature number) is the learnable weight matrix. The representation of input graph $\mathcal{G}$ and negative graph $\tilde{\mathcal{G}}$ are obtained by the GCN-based encoder, i.e. $\mathbf{H} = f(\mathbf{X}, \mathbf{A}), \tilde{\mathbf{H}} = f(\tilde{\mathbf{X}}, \tilde{\mathbf{A}})$.

DGI considers a noise-contrastive type objective with a standard binary cross-entropy (BCE) [16] loss between the samples from the positive view and the negative view. The loss function is defined as

$$\mathcal{L} = \frac{1}{n+m}\left(\sum_{i=1}^{n}\mathbb{E}_{(\mathbf{X},\mathbf{A})}\left[\log D\left(\vec{h}_i, \vec{s}\right)\right] + \sum_{j=1}^{m}\mathbb{E}_{(\tilde{\mathbf{X}},\tilde{\mathbf{A}})}\left[\log\left(1 - D\left(\vec{\tilde{h}}_j, \vec{s}\right)\right)\right]\right), \tag{6}$$

where $\vec{h}_i$ and $\vec{\tilde{h}}_j$ are the $i$-th spot of $\mathbf{H}$ and $j$-th spot of $\tilde{\mathbf{H}}$, respectively. $\vec{s}$ is high level representation, obtained by readout function $R(\cdot)$. $D(\cdot,\cdot)$ is defined as a discriminator, which outputs a probability score.

For the readout function $R(\cdot)$, DGI calculates a simple average of the representations for all nodes

$$R(\mathbf{H}) = \sigma\left(\frac{1}{n}\sum_{i=1}^{n}\vec{h}_i\right), \tag{7}$$

where $\sigma(\cdot)$ denotes the logisitic sigmoid nonlinearity. Here, the high-level representation obtained by the readout function is the high-level summaries of the graph, which contain global information.

For the discriminator $D(\cdot,\cdot)$, DGI scores the probability using a simple bilinear scoring function

$$D\left(\vec{h}_i, \vec{s}\right) = \sigma\left(\vec{h}_i^T \boldsymbol{\Theta}\vec{s}\right), \tag{8}$$

where $\boldsymbol{\Theta} \in \mathbb{R}^{d' \times d'}$ is a learnable scoring matrix and $\sigma(\cdot)$ is the logisitic sigmoid nonlinearity. Here, $D(\vec{h}_i, \vec{s})$ represents the consistency between representation of node $v_i$ and graph level representation $\vec{s}$. Meanwhile, $1 - D(\vec{\tilde{h}}_j, \vec{s})$ represents the inconsistency between the representation of node $\tilde{v}_j$ for negative view and the graph level representation $\vec{s}$.

GRAS4T compares the two positive views with the original view and the negative view, respectively, and then the discriminator $D$ needs to be redefined. Taking local-glocal contrastive loss as an example, GRAS4T computes the relationship between high-level representations $\vec{s}_1, \vec{s}_2$ and representation $\vec{h}_i$ by utilizing the discriminant $D(\cdot,\cdot,\cdot)$, which can be formulated as

$$D\left(\vec{h}_i, \vec{s}_1, \vec{s}_2\right) = \sigma\left(\vec{h}_i^T \boldsymbol{\Theta}\vec{s}_1 + \vec{h}_i^T \boldsymbol{\Theta}\vec{s}_2\right). \tag{9}$$

#### 2.2 subspace analysis

Given a set of samples $\mathbf{X} = [\mathbf{X}_1, \cdots, \mathbf{X}_k] = [\boldsymbol{x}_1, \boldsymbol{x}_2, \cdots, \boldsymbol{x}_n] \in \mathbb{R}^{d \times n}$ drawn from an unknown union of $k$ subspaces $\{S_i\}_{i=1}^{k}$ of unknown dimensions $d_i = dim(S_i), 0 < d_i < d, i = 1, \cdots, k$ and $\mathbf{X}_i$ is a set of samples from subspace $S_i$ with sample size $n_i$, where $n = \sum_{i=1}^{k} n_i$ [17]. Based on the assumption of subspace analysis, each data point $x_i$ is a linear combination of other data points in the same subspace, implying that the self-expressive matrix should be block diagonal in theory. Further, the self-expressive model is a crucial component of subspace analysis [18]. More specifically, the key problem in subspace analysis involves obtaining the self-expressive matrix $\mathbf{C}$ by solving the following optimization problem

$$\min_{C} L(\mathbf{XC}, \mathbf{X}) + \beta \left\|\mathbf{C}\right\|_{\xi}, \; s.t. \; diag(\mathbf{C}) = 0, \tag{10}$$

where $\mathbf{X}$ is the dataset and $\beta$ is the trade-off parameter. The $diag(\mathbf{C}) = 0$ means that the diagonal of $\mathbf{C}$ is restricted to 0, which is used to prevent trivial solutions. The first term of the objective function reconstructs each sample from all other samples, indicating the difference between the input data and the represented data. The second term introduces a regularisation of the coefficient matrix. Based on a priori information, different norms $\xi$ (including $l_p$-normalization, Frobenius-normalization, and nuclear norm regularizer) are used to obtain a self-expressive matrix with different reconstruction properties, such as sparsity, low-rank, and connectivity.

Due to the complexity of data in the real world, an increasing number of models are designed depending on the background of the problem and the assumptions about the data distribution [19, 20, 21]. Efficient kernel graph convolutional subspace clustering (EKGCSC) is a subspace clustering method introduced in [22]. EKGCSC considers the rich spatial structure information of the data and the non-linear relationships between points, the model seeks a block diagonal representation by minimization

$$\min_{\mathbf{C}} \frac{1}{2} \left\| \Phi(\mathbf{X}) \hat{\mathbf{D}}^{-\frac{1}{2}} \hat{\mathbf{A}} \hat{\mathbf{D}}^{-\frac{1}{2}} \mathbf{C} - \Phi(\mathbf{X}) \right\|_F^2 + \frac{\beta}{2} \left\| \mathbf{C} \right\|_F^2 . \tag{11}$$

Here, $\|\mathbf{C}\|_F$ denotes the Frobenius norm of matrix $\mathbf{C}$. $\Phi : \mathbb{R}^m \to \mathcal{H}$ is a mapping from the input space to the reproducing kernel Hilbert space $\mathcal{H}$. The elements in the kernel Gram matrix $\mathbf{K_{XX}}$ consist of the inner product of vectors in the Hilbert space $\mathcal{H}$, i.e. $[\mathbf{K_{XX}}]_{ij} = [\langle \Phi(\mathbf{X}), \Phi(\mathbf{X}) \rangle_{\mathcal{H}}] = \Phi(x_i)^T \Phi(x_j) = k(x_i, x_j)$. $\beta$ is a balance parameter. KGCSC set Gaussian kernel as kernel function $k$, i.e. $k(x_i, x_j) = \exp\left(-\gamma \left\| x_i - x_j \right\|^2\right)$. The above objective function has a closed-form solution and the detail of the optimization process is provided in [22].

#### 3 Additional Experimental Results

Here, we present additional experimental results. Figure S1 shows the ARI and NMI scores of the six spatial domain identification methods on the DLPFC dataset. Figure S2 displays the results of the ablation study on the DLPFC dataset. Figure S3 depicts the UMAP and PAGA visualization results of GRAS4T on the DLPFC dataset. Figure S4 shows the spatial domains of human HER2-positive breast tumor (HER2+) detected by STAGATE, CCST, and GRAS4T. Figure S5 demonstrates the NMI scores of the spatial domain identification results on the DLPFC and HER2+ datasets. Figure S6 shows the spatial domain identification results of the six methods on the human breast cancer dataset. Figure S7 displays the heatmap of inter-domain Pearson correlation coefficients for the four methods on the 151672 slice of the DLPFC dataset and the A1 slice of the HER2+ dataset. Figure S8 shows the spatial domain identification results in the mouse brain anterior&posterior dataset detected by STAGATE, CCST, and GRAS4T. Figure S9 shows the spatial domain identification results on 12 slices of the mouse hypothalamus datasets detected by STAGATE, CCST, and GRAS4T. Table S4 demonstrates the closeness of the three spatial domain identification methods to the reference cell type on the mouse hypothalamus datasets.

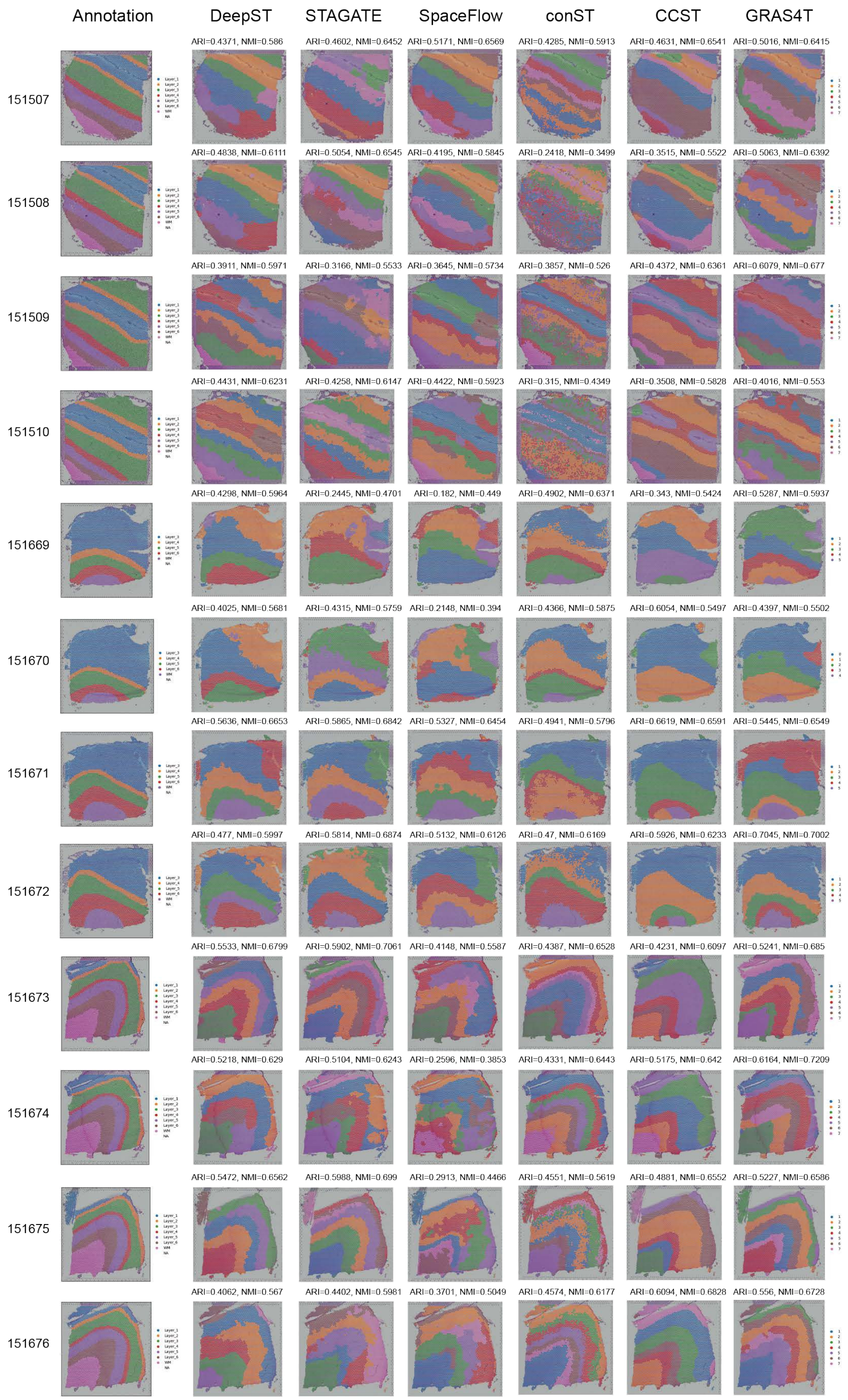


Figure S1: Manual annotations and comparison of spatial domains identified by DeepST, STAGATE, SpaceFlow, conST, CCST, and GRAS4T on the 12 slices of DLPFC dataset.

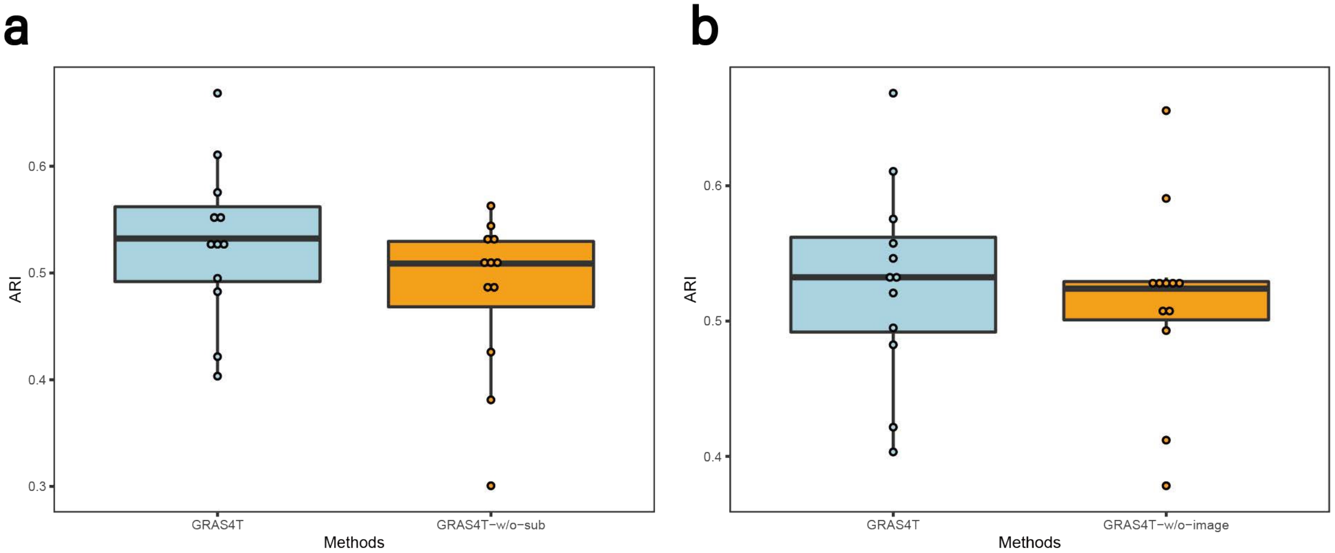


Figure S2: Ablation study of different measured by ARI. (a) Comparison of ari scores on DLPFC between GRAST and GRAS4T without subspace module. (b) Comparison of ari scores on DLPFC between GRAST and GRAS4T without H&S image-based augmentation.

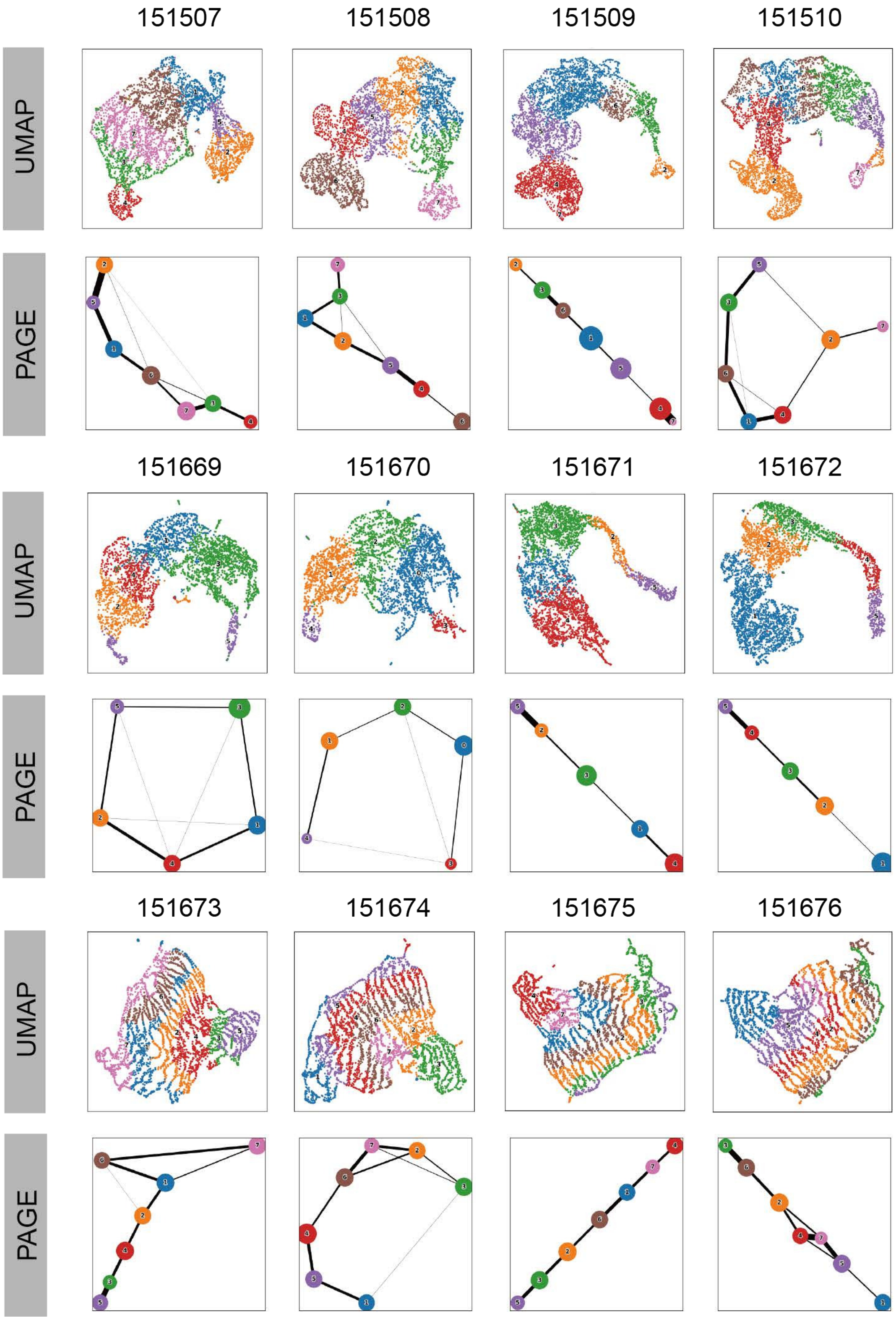


Figure S3: UMAP visualization and PAGA graphs generated by GRAS4T on the 12 slices of DLPFC dataset.

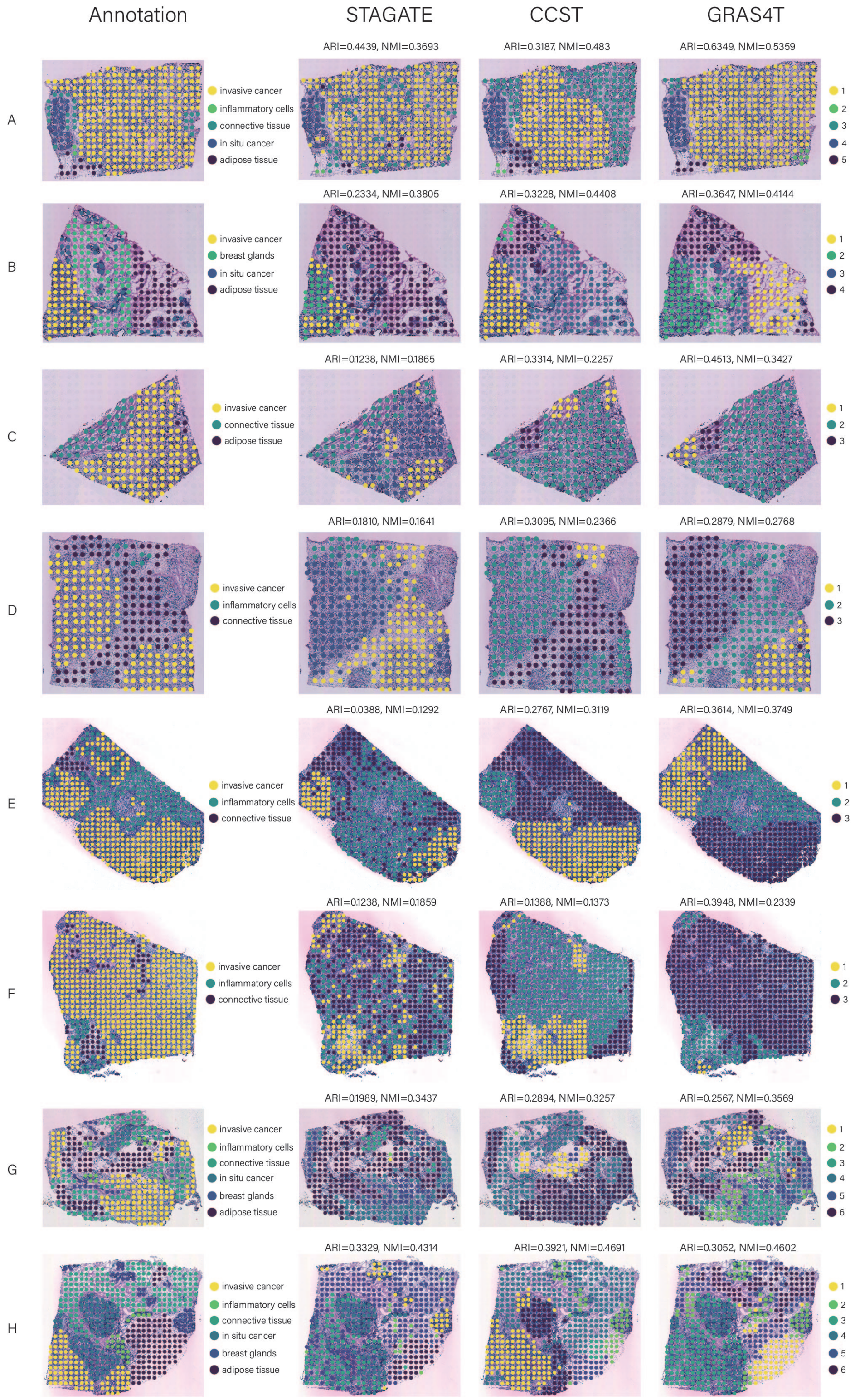


Figure S4: Manual annotations and comparison of spatial domains identified by STAGATE, CCST, and GRAS4T on the 8 slices of HER2+ dataset.

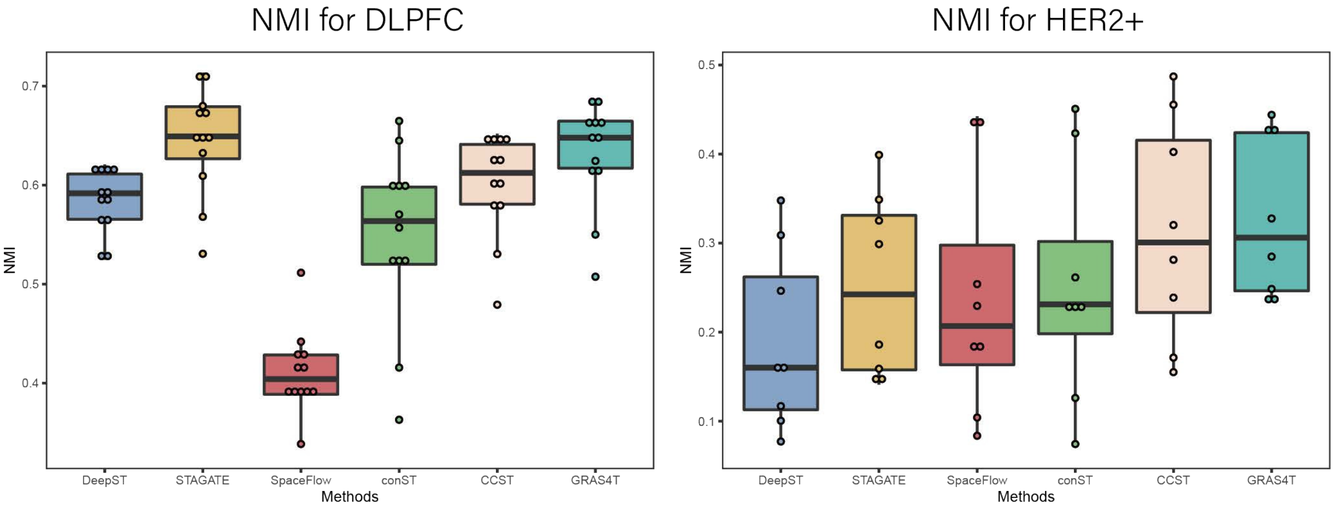


Figure S5: Spatial domain identification performances of six methods in terms of NMI. (a) Boxplot of clustering accuracy in all sections of the DLPFC dataset in terms of NMI values for all methods. (b) Boxplot of clustering accuracy in all sections of the HER2+ dataset in terms of NMI values for all methods.

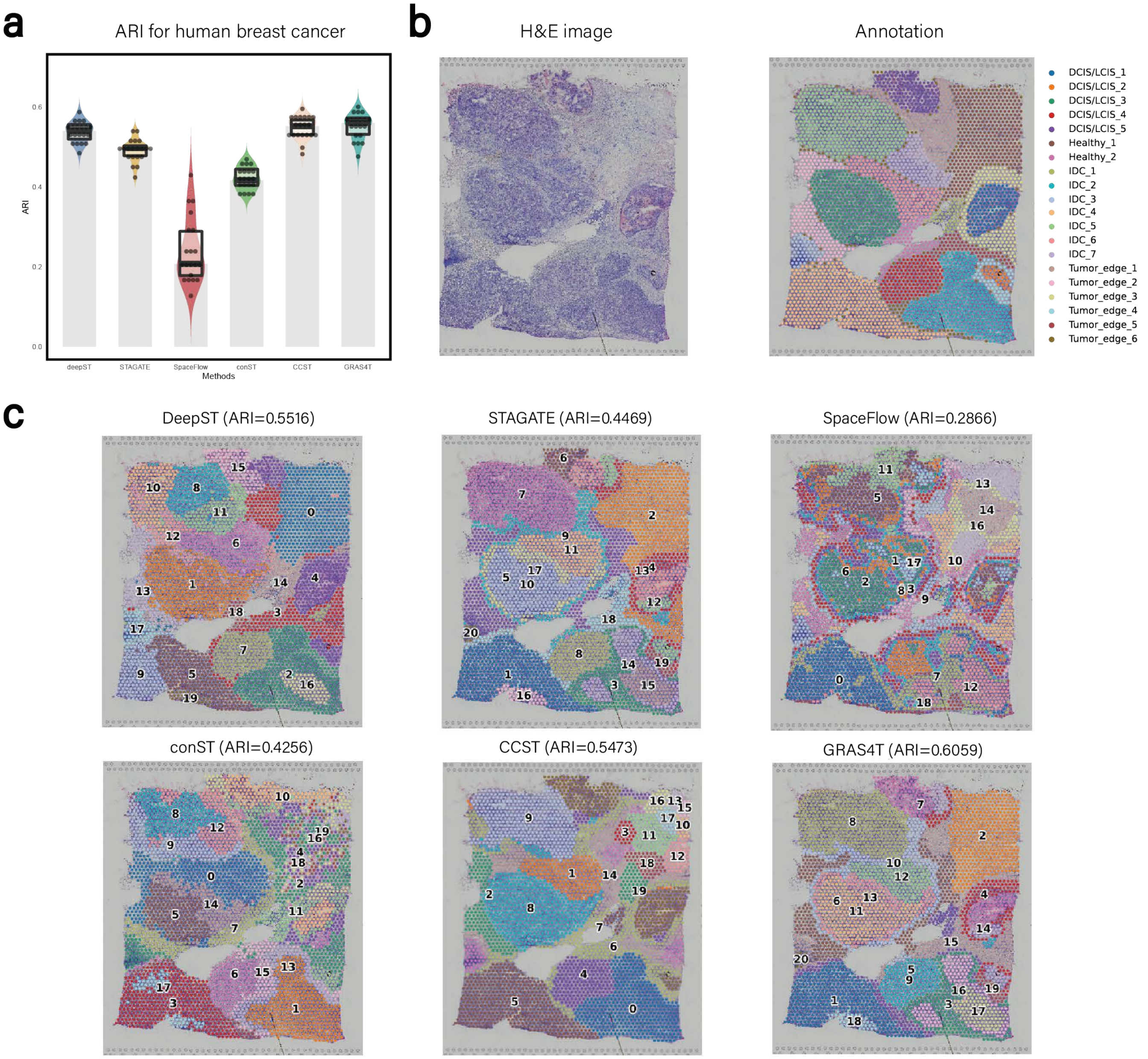


Figure S6: Comparison of spatial domains identified by DeepST, STAGATE, SpaceFlow, conST, CCST, and GRAS4T in the human breast cancer dataset. (a) Comparison of different methods by ARI pirate graph. (b) H&E image and manual annotation. (c)Visualization of spatial domains in (b) identified by six methods.

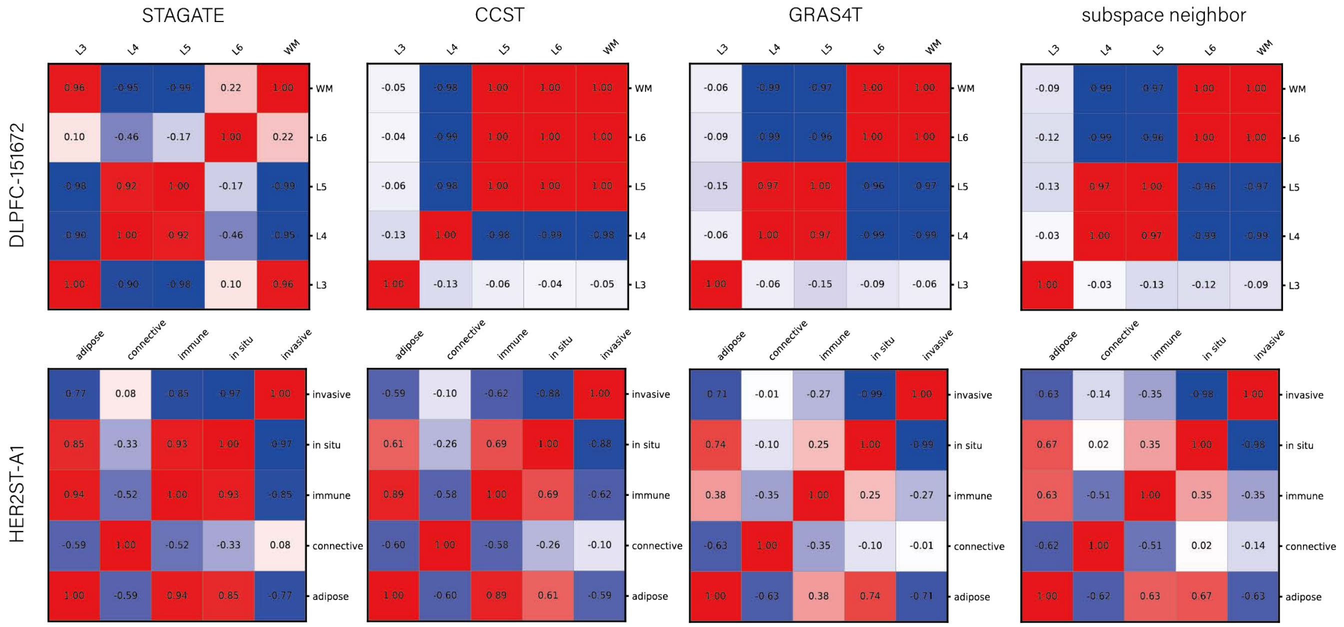


Figure S7: Heatmap of inter-domain Pearson correlation coefficients for STAGATE, CCST, GRAS4T, and GRAS4T using subspace nearest neighbor adjustment.

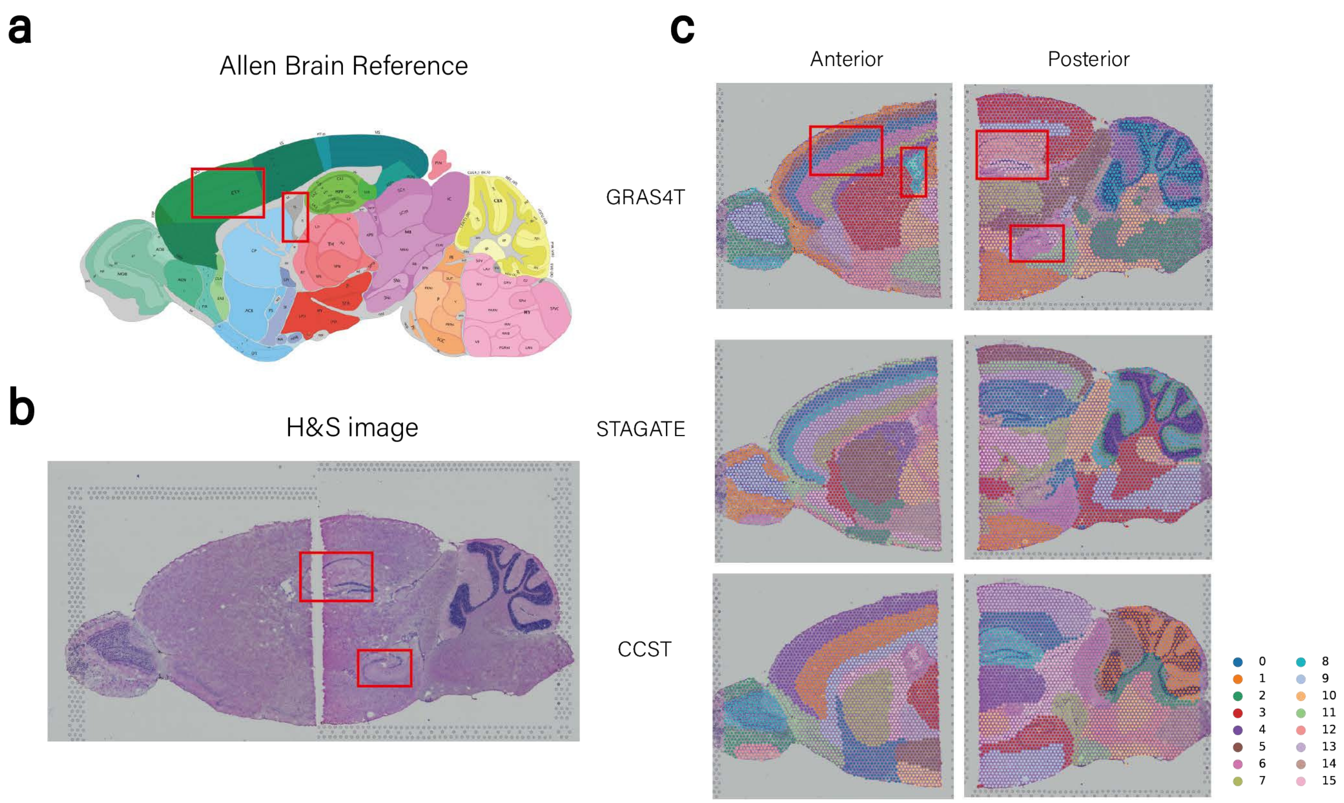


Figure S8: Spatial domain identification in the mouse brain anterior&posterior dataset. (a) Annotated brain section image from Allen Mouse Brain Atlas for reference. (b) H&E image of mouse brain anterior and posterior. (c) Spatial domains in (b) detected by GRAS4T, STAGATE, and CCST.

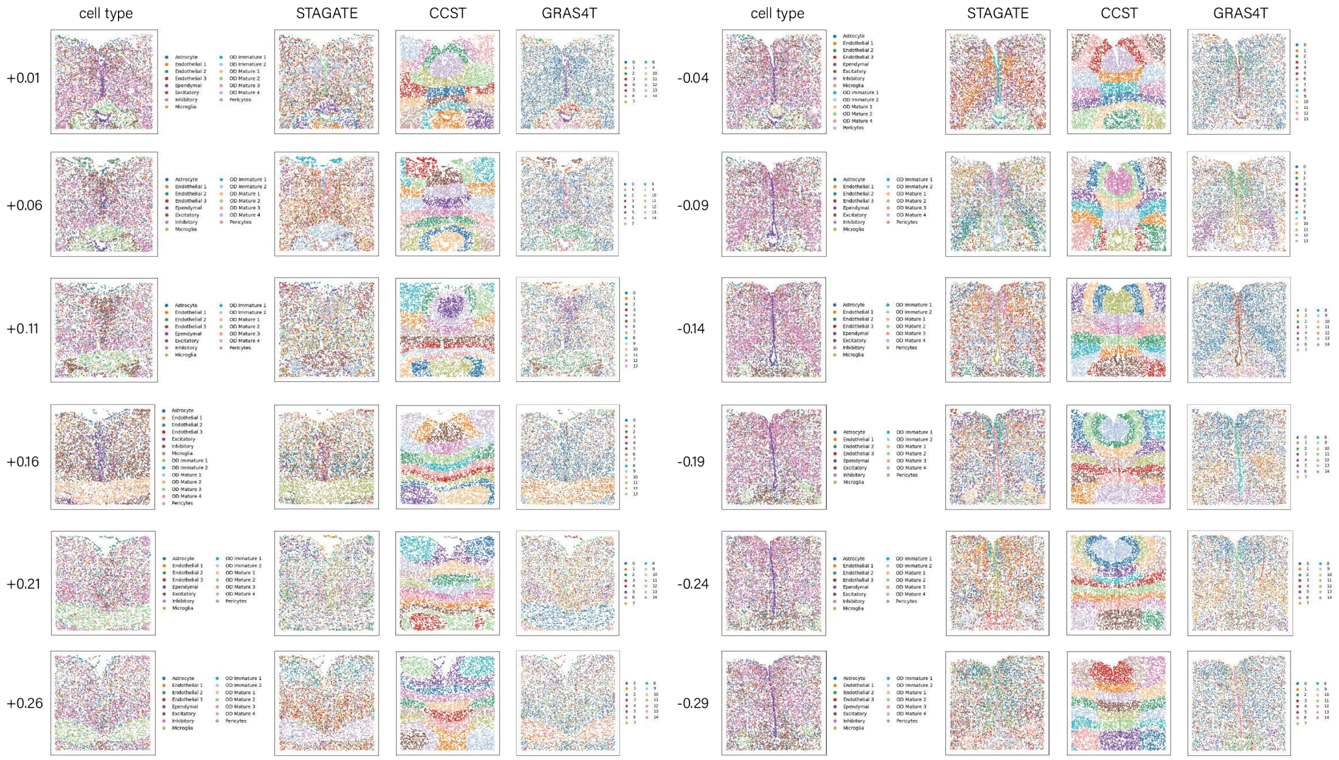


Figure S9: Cell type and spatial domain identification by STAGATE, CCST, and GRAS4T on the 12 slices of mouse hypothalamus datasets.

Table S4: The closeness (Using ARI metrics) of the three spatial domain identification methods to the reference cell type on the mouse hypothalamus datasets.

| slice | STAGATE | CCST | GRAS4T |
|---|---|---|---|
| +0.01 | 0.0826 | 0.0317 | 0.5005 |
| +0.06 | 0.0682 | 0.0455 | 0.4418 |
| +0.11 | 0.1078 | 0.0526 | 0.4495 |
| +0.16 | 0.0719 | 0.0494 | 0.4641 |
| +0.21 | 0.0264 | 0.0443 | 0.4206 |
| +0.26 | 0.0451 | 0.0186 | 0.3780 |
| -0.04 | 0.0805 | 0.0175 | 0.3818 |
| -0.09 | 0.0403 | 0.0249 | 0.2784 |
| -0.14 | 0.0754 | 0.0257 | 0.4711 |
| -0.19 | 0.0708 | 0.0236 | 0.3970 |
| -0.24 | 0.1530 | 0.0333 | 0.3863 |
| -0.29 | 0.1167 | 0.0251 | 0.4284 |

**14.** Yixin Liu, Ming Jin, Shirui Pan, Chuan Zhou, Yu Zheng, Feng Xia, and Philip Yu. Graph self-supervised learning: A survey. *IEEE Transactions on Knowledge and Data Engineering*, 2022.
